# A meta-analysis of task-based differences in bilingual L1 and L2 language networks

**DOI:** 10.1101/2021.12.28.474335

**Authors:** Lindy Comstock, Bruce Oliver

## Abstract

The functional organization of first (L1) and second (L2) language processing in bilinguals remains a topic of great interest to the neurolinguistics community. Functional magnetic resonance imaging (fMRI) studies report meaningful differences in the location and extent of hemodynamic changes between tasks performed in the L1 and L2, yet there is no consensus on whether these networks can be considered truly distinct. In part, this may be due to the multiplicity of task designs implemented in such studies, which complicates the interpretation of their findings. This paper compares the results of previous bilingual meta-analyses to a new ALE meta-analysis that categorizes neuroimaging studies by task design. Factors such as the age of L2 acquisition (AoA) and the L2 language proficiency level of participants are also considered. The findings support previous accounts of the effect of participant characteristics on linguistic processing, while at the same time revealing dissociable differences in fMRI activation for L1 and L2 networks within and across tasks that appear independent of these external factors.

## 1. Introduction

The neurobiology and functional organization of L1 and L2 processing continue to elude researchers, with neuroimaging studies consistently yielding conflicting results for the localization of each language and the extent to which their neural representations overlap. This is particularly true when attempting to differentiate specific language competencies within each network. In recent years, bilingualism researchers have instead turned their attention to characterization of the language control network in bilinguals (Hernández, Costa, Fuentes, Vivas, & Sebastián-Gallés, 2010; Pliatsikas, & Luk, 2016; Rodriguez-Fornells, De Diego Balaguer, & Münte, 2006), structural magnetic resonance imaging (MRI) studies (Abutalebi, Canini, Della Rosa, Green, & Weekes, 2015; Burgaleta, Sanjuán, Ventura-Campos, Sebastian-Galles, & Ávila, 2016; Kuhl, Stevenson, Corrigan, van den Bosch, Can, & Richards, 2016), and connectivity research (Berken, Chai, Chen, Gracco, & Klein, 2016; Sulpizio, Del Maschio, Del Mauro, Fedeli, & Abutalebi, 2020; Zou, Abutalebi, Zinszer, Yan, Shu, Peng, & Ding, 2012). While these studies foreground important research questions in their own right, the shift in the literature reflects a growing frustration with our abilities to discern potentially fine-grained differences between L1 and L2 processing by means of univariate fMRI analyses. Moreover, numerous factors that are thought to be relevant to linguistic processing—such as AoA, proficiency level, the linguistic distance between L1 and L2, and frequency of language use—may vary considerably among bilinguals. The complexity of the research population therefore complicates our ability to establish general conclusions that would characterize the population as a whole.

Instead, it has become commonplace to assume that multiple languages map onto essentially the same neuroanatomy, utilized in subtly different ways. However, this assumption appears to be driven more by the inconsistencies observed in individual fMRI studies and their inability to substantiate a comprehensive model of bilingual language processing, rather than any conclusive findings. In fact, evidence from methodologies with greater temporal precision (electroencephalography and event related potentials: Barac, Moreno, & Bialystok, 2016; Heidlmayr, Hemforth, Moutier, & Isel, 2015; Morales, Yudes, Gómez-Ariza, & Bajo, 2015; Paap, Sawi, Dalibar, Darrow, & Johnson, 2015; Zhang, Kang, Wu, Ma, & Guo, 2015, and magnetoencephalography: Wang, Xiang, Vannest, Holroyd, Narmoneva, Horn, Liu, Rose, deGrauw, & Holland, 2011) or with greater spatial precision (electrocortical stimulation mapping: Fernández-Coello, Havas, Juncadella, Sierpowska, Rodríguez-Fornells, & Gabarrós, 2016, and intercranial electrocorticography: Cervenka, Boatman-Reich, Ward, Franaszczuk, & Crone, 2011) suggest that L1 and L2 processing may diverge more substantively than can be currently be established. Therefore, the possibility of constructing a coherent account of differences between the two networks should not be disregarded.

One method to increase the statistical power and generalizability of fMRI findings is to conduct a meta-analysis that operates on the combined results of multiple studies. If differences in L1 and L2 processing are present but too subtle to detect in fMRI studies with limited power, such an approach might enable the identification of individual networks. Today, papers investigating L1 and L2 language processing in bilinguals number in the hundreds. This has allowed for no less than six neuroimaging meta-analyses devoted to various aspects of bilingual language use (Cargnelutti, Tomasino, & Fabbro, 2019; Indefrey, 2006; Liu & Cao, 2016; Luk, Green, Abutalebi, & Grady, 2012; Sebastian, Laird & Kiran, 2011; Sulpizio, Del Maschio, Fedeli, & Abutalebi, 2020). However, the specific nature of many research questions in the field has facilitated experimental paradigms that defy easy categorization or comparison, frequently leading to the combination of data types that are not compatible in effort to increase the size of the overall data set, which in turn weakens the validity of the summarized findings. These differences in data type include task design, participant background, and analysis strategy. Additional complications that may be less readily apparent arise from the absence of a common method to define and measure descriptive categories such as proficiency level, degree of language exposure, and AoA, a failure to report these categories, and the choice of which data contrasts to include into the meta-analysis.

The existing meta-analyses adopt different strategies for inclusion criteria when faced with heterogeneous data: some average across task design in attempt to group studies by broad linguistic domains such as syntax, semantics, and so forth (Indefrey, 2006; Sulpizio, Del Maschio, Fedeli, & Abutalebi, 2020, others abandon the attempt to differentiate linguistic domains in favor of participant background characteristics like AOA or proficiency (Cargnelutti, Tomasino, & Fabbro, 2019; Sebastian, Laird & Kiran, 2011), and one study estimates the degree of similarity between task designs such that all linguistic competencies are claimed to be represented equally within L1 and L2 conditions (Liu & Cao, 2016). Additional choices the authors must make include what contrasts or categories to analyze, which may include a linguistic task performed in each language separately or both combined, and the control condition, which may rest or a fixation cross or a task carefully matched to the experimental one. Some studies compare the L1 in bilinguals to monolinguals, or the L1 or L2 in one set of bilinguals to a second set. These choices about how to compile a data set are often not explicitly mentioned and occasionally misrepresented. Combining data with different underlying assumptions lessens the chance that the results will converge across studies and may overrepresent data of one underlying type. This may explain why existing meta-analysis show conflicting findings and differences in L1 and L2 processing remain unreported.

This paper first establishes the need for a task-based meta-analysis, and then reviews the existing neuroimaging meta-analyses devoted to bilingual language processing. Finally, we present our own ALE meta-analysis in which we investigate how grouping studies by a common experimental paradigm may inform the bilingual neuroimaging literature about which brain regions can be associated with L1 and L2 processing.

### 1.1 Why a Task-Based Meta-analysis?

Nearly fifteen years ago, the first meta-analysis of bilingual neuroimaging studies (Indefrey, 2006) took a close look at task design in attempt to control for the various linguistic processing levels required by individual tasks. The author noted a trend in the monolingual neuroimaging literature away from simpler methods towards more complex paradigms and predicted a similar trajectory for bilingual studies. At the same time, Indefrey expressed concern that the variability in experimental design details he observed could potentially obscure differences in L1 and L2 processing, given our limited knowledge about how manipulating design features may affect an analysis.

As a result, Indefrey (2006) organized studies according to the principle of whether their tasks included certain core components of language processing, such as lemma or word form retrieval, graphemic or phonological word representations, syllabification, phonetic encoding, shared “lead-in processes” (Indefrey & Levelt, 2000, 2004), and so forth. This method goes beyond the broader and somewhat simplistic parcellation of studies by “linguistic domain” (syntax, semantics, phonology, etc.) adopted by subsequent meta-analyses. The assumption that different tasks can be combined into the same category for analysis is founded on the belief that they will be processed identically by the neural substrate. Despite Indefrey’s (2006) attempt to control for the linguistic subcomponents shared between tasks, his analysis nonetheless necessitated the inclusion of widely differently experimental paradigms that introduced variation in non-linguistic processing factors. To date, no other bilingual meta-analysis has attempted to classify studies by task design with the same rigorous criteria.

The clinical language mapping literature has independently raised concerns about the assumption that task design can be conflated with underlying cognitive processes. Ivanova, Dragoy, Kuptsova, Akinina, Petrushevskii, Fedina, Turken, Shklovsky, & Dronkers (2018) draw attention to the widespread inconsistencies found in where studies localize verbal working memory within and across hemispheres. In fact, in comparing two common measures of verbal working memory, Ivanova et al. (2008) were able to entirely disassociate the two tasks, obtaining maps that shared no common anatomical areas or voxels. Similar to Indefrey (2006), the authors question the variety of tasks implemented in verbal work memory research and the fact that different tasks are assumed to be interchangeable. The authors attribute inconsistencies in the wider literature to a failure to consider what subprocesses are recruited for each task. It may be that the inconsistent localization of L1 and L2 linguistic processes observed in the bilingual neuroimaging literature stems from the same considerations.

However, even an approach based on careful consideration of the component subprocesses involved in a linguistic task of interest can fall victim to invalid assumptions about how to define tasks, their subcomponents, and their effect on fMRI activation (Bookheimer, Zeffiro, Blaxton, Gaillard, & Theodore, 1995). For example, the difference in tasks as similar as reading aloud or reading covertly cannot be distilled to the simple addition of motor activity. Adding or changing a subprocess may substantially influence BOLD activity, disrupting the pattern of activation previously observed. Overt reading tasks have in fact been shown to yield less fMRI activation in traditional language areas (Bookheimer et al., 1995). Moreover, research from the monolingual language processing literature suggests that no linguistic processing level can be related to a single cortical area (Indefrey & Cutler, 2004), highlighting the potential for distributed rather than strictly modular language networks.

The clinical literature has also found discrepancies in obtaining the same extent, location, and lateralization of fMRI activation between studies that utilize active and passive control conditions (Binder, Swanson, Hammeke, & Sabsevitz, 2008). This important design feature is neglected when meta-analyses combine different contrast types in the same analysis. An active baseline would be a task that closely matches the experimental condition, with the aim of isolating the linguistic element of the experimental task from the other sensory or cognitive processes involved. A passive baseline is typically rest or a very minimal task such as viewing a fixation cross. Binder et al. (2008) found that passive baselines produced wider and more varied fMRI activation, whereas active controls may produce erroneous findings: it is possible for greater deactivation in the baseline condition to drive an effect, rather than true activation in the experimental task. Two bilingual meta-analyses have addressed the use of active baselines (Indefrey, 2006; Luk et al. 2012) to different effect, but an in-depth discussion of their potential to generate erroneous findings has thus far been absent in the bilingual neuroimaging literature.

Finally, it is well known that bilinguals use their languages selectively across tasks and linguistic domains (Francis, 2004; Grosjean, 2008, 2012). One language is often restricted to use in the home or informal environments, while the other may be required for professional discourse or formal occasions (Hoff, 2006; Polinsky, & Kagan, 2007). A large-scale study investigating the bilingual advantage in executive control tasks has shown that the variable strength of findings across such studies may be explained by numerous social, contextual, and performance factors (Van den Noort, Struys, Bosch, Jaswetz, Perriard, Yeo, & Lim, 2019). This is likely to hold true for standard linguistic tasks as well. It has been suggested that the relative importance of the frequency of language use may outweigh that of proficiency (Verreyt, Woumans, Vandelanotte, Szmalec, & Duyck, 2016). Thus, the degree to which each language is typically recruited for the linguistic structures employed in an experimental task may vary considerably and affect the overall result in a neuroimaging study.

Complex syntax and reading tasks serve as examples of this, as they are more likely to be performed in a formal environment. In fact, highly educated individuals have been shown to place greater weight upon syntactic cues in judgment tasks (Chipere, 1998; Dabrowska, 1997; Schumann, 2007). Thus, bilinguals who are weaker in their L2 proficiency overall but utilize the L2 in formal environments may be selectively stronger in their L2 for syntactic or reading tasks, while remaining stronger in their L1 for picture naming or passive listening. To our knowledge, individual differences in relative L1/L2 proficiency by task has not been addressed by the bilingual neuroimaging literature and therefore is beyond the scope of a meta-analysis. However, the bilinguals who participate in research experiments are often university students who have learned their L2 for instrumental purposes that require mastery of formal language, and the formality level of a task is likely to be assessed similarly by all participants. Therefore, differentiating tasks may partially control for processing strategies related to formal and informal language use.

Numerous reasons support the rationale to organize a meta-analysis around task design. The present study takes a maximally conservative approach: because the complexity of processing demands in linguistic tasks means we cannot account for all of them, only studies that utilize a common task design will be included. Studies with active and passive baselines will be accepted; however, they will be controlled for in another fashion: only data from the L1>L2 or L2>L1 contrasts will be entered into the meta-analysis. This means that extraneous processing demands should be eliminated, because the contrast is performed between the experimental task when conducted in the L1 and the L2, rather than pooling the results of a contrast performed between the task and a baseline condition in each language separately. While a conservative approach may restrict the greatest possible extent of fMRI activations, the results that are obtained can be considered with a high degree of confidence.

### 1.2 A Review of Previous Meta-analyses

<u>Indefrey (2006)</u> presents an early review of bilingual neuroimaging studies that acknowledges the importance of task design, linguistic processing level, and control conditions for the interpretation of complex results. The meta-analysis includes 30 experiments from 24 studies and employs a statistical method that was first developed by the author to study language processing in monolinguals (Indefrey, 2004; Indefrey & Cutler, 2004; Indefrey & Levelt, 2000, 2004). This method calculates the chance level that a region will be reported within a descriptive reference system of 114 regions across the whole brain (Indefrey & Levelt, 2000, 2004). To reach significance, the chance probability must be less than 0.5 that the same foci will be found in each individual experiment. Only differences between the L1 and L2 networks are reported, which suggests that the L1>L2 and L2>L1 contrasts were used, as does the small number of brain regions identified. However, the data type is not explicitly stated.

A whole brain fMRI analysis encompasses a huge number of voxels—a three-dimensional unit of brain tissue. Therefore, calculations must be corrected for the numerous comparisons that are made in a whole brain statistical analysis. Region of interest (ROI) studies consider fMRI activation only within a limited, pre-determined area of the brain, such that the number of comparisons is reduced, and the statistical threshold for determining the significance level is decreased. Indefrey’s (2006) method, unlike those employed in later meta-analyses, allows for whole brain and ROI studies to be compared on equal footing, and for this reason, the author legitimately includes ROI studies that are absent from later meta-analyses. However, his method does not correct for the fact that ROI analyses may be constrained by prior assumptions about where activation should be found.

Neuroimaging studies were divided into categories of tasks that require the same linguistic subprocesses, according to their task design. The resulting data set included 5 to 6 experiments per task (word generation, picture naming, semantic decision) and 14 experiments for a combined task (sentence-level listening and reading). Given that the number of studies, and therefore, the number of reported regions were low, the chance probability level was also low. Clusters had only to be reported in two or three studies, based on category size, to pass the reliability criterion. The relative influence of each study will be a primary concern in all of the meta-analyses we consider.

The **semantic decision** task elicited widely dispersed clusters and was the only task to engage posterior and inner cortical regions and regions in **both networks**. The category combined association and decision tasks requiring a metalinguistic judgment and button press. The results were generated by three of six experiments from two studies. The two studies that produced no results for the L2 network instead identified clusters in the L1 network, which suggests considerable heterogeneity in the data.

Word level comprehension studies comprised **word generation** and **picture naming.** Only one region was identified in the **L2 network—** the left *inferior frontal gyrus*, with bilateral results for word generation. Only two of five experiments in each task contributed to these findings.

**Sentence-level listening and reading** tasks generated clusters in the **L2 network** in only six of fourteen studies. These results were driven by bilinguals with high L2 proficiency and exposure, and most often in tasks that required a button press. Individual studies reported findings in the L1 network, but no region replicated in more than one study.

Overall, the **L2 network** was concentrated in left frontal regions of the classical language network. This was true for active tasks, whereas passive tasks generated increased fMRI activation only upon addition of an explicit metalinguistic decision component. Elsewhere, passive tasks have been shown to elicit less hemodynamic response (Binder et al., 2008). Yet the metalinguistic component and button press are unlikely to underlie the activation found in the **L1 network,** as both typically engage frontal regions (Müller & Basho, 2004).

Indefrey (2006) remains optimistic that reliable disassociations between L1 and L2 networks can be found if both participant background characteristics and task type are considered. He concludes that three factors (late AoA, low proficiency level, low degree of exposure) may target different components of linguistic processing: word-level production appears to be influenced by all three factors, word-level comprehension is influenced by proficiency level in semantic decision tasks, and sentence comprehension tasks that include a decision component are influenced by AoA. Furthermore, the author speculates that L2 may activate non-lexical compositional, syntactic, or decision processes in the left *inferior frontal gyrus* more strongly than it does lexical process in the *temporal lobe*, particularly for bilinguals with late AoA or low L2 proficiency. While still preliminary in scope, Indefrey (2006) provides a number of compelling working hypotheses. The question remains whether the sparse regions identified here reflect the selection criteria, the underlying data set, or a lack of sensitivity in the method.

<u>Sebastian, Laird, & Kiran (2011)</u> conducted a meta-analysis of bilingual neuroimaging studies that focuses on the effect of L2 proficiency level, which they consider to be most relevant among potential mediating factors. L1 and L2 networks were investigated in separate conjunction analyses for participants with high and low/moderate proficiency, yielding four sets of results. At the same time, the authors acknowledged difficulty in differentiating groups due to the heterogeneity of language proficiency scales adopted by study authors. Several studies utilized by Indefrey (2006) were excluded as ROI studies or because they did not report the participant proficiency level or stereotactic coordinates. Therefore, although conducted several years later, this meta-analysis comprised a smaller data set: 14 published studies with 17 experiments (11 with high and 6 with low proficiency participants).

This meta-analysis was the first to implement the activation likelihood estimation (ALE) technique for bilingual neuroimaging studies (Eickhoff et al., 2009; Laird, Fox, et al., 2005; Turkeltaub, Eden, Jones, & Zeffiro, 2002). ALE is a statistical method that computes the probability of activation within each voxel for a given contrast based on the spatial distribution of the foci reported in each study. The method performs a random-effects analysis by calculating the above-chance clustering between experiments. Author-supplied anatomical labels can be avoided, rendering an ALE analysis more precise when comparing across meta-analyses; however, it requires the exclusion of ROI studies and those without published coordinates. All subsequent bilingual meta-analyses use this method. Calculations were corrected for multiple comparisons using the FDR algorithm (Genovese, Laxar, & Nochols, 2002) with a somewhat lenient significance threshold of *p* < .05 and a minimum cluster size of 150 mm^3^.

Sebastian et al. (2011) analyzed data from contrasts comparing L1 or L2 relative a baseline condition. This decision, together with the low significance threshold, may explain why relatively large clusters were obtained. Tasks involving suprasegmental processing, language switching, or translation were excluded, as well as ‘between subjects’ studies. The authors justified combining a wide range of heterogeneous tasks in their data set, citing an argument found in Hagoort (2005) and Bookheimer (2002) that the left inferior frontal gyrus does not play a modality or content specific role in linguistic processing, and extending that argument broadly to all regions within the language network. Upon consideration of the tasks found in each subset for analysis, it is notable that they tend to be specific to each proficiency level: only semantic judgment and passive listening tasks occur in both the high and low/moderate proficiency group.

The tasks performed by less proficient bilinguals involve more processing levels and greater task complexity: word reading, sentence generation and comprehension, and syntactic judgment. Highly proficient bilinguals performed picture naming, noun/verb generation, and homophone mapping. Three out of six tasks in the less proficient group involve the sentence and discourse level, whereas the high proficiency group contained only one task out of five with stimuli above the word level. The authors note that clusters obtained from the less proficient group were smaller and distributed widely over both hemispheres. However, this clustering pattern is also consistent with the results of an analysis using complex tasks.

In the **high proficiency group**, L1 and L2 networks overlapped considerably and were distributed across left hemisphere regions, with larger clusters in frontal than in posterior regions. Contradicting Indefrey’s (2006) findings, the L1 network was more extensive. In the **low proficiency group**, the L1 network was left-lateralized, but with more right hemisphere and bilateral clusters, particularly for the L2. **Both networks** center around known language areas, but right hemisphere clusters appear in regions not considered to be language specific. Clusters specific to the **L1 network** are more posterior, whereas the **L2 network** recruited widespread frontal regions.

Sebastian et al. (2011) note that three of their four meta-analyses yielded highly similar results. However, the L2 network in the low proficiency group appeared to differ in several respects. Clusters were smaller with a wider distribution across both hemispheres, which the authors interpreted as the result of the increased processing demands associated with a less proficient language. The *anterior cingulate* and *paracingulate gyrus* were uniquely activated for this group, in addition to the *dorsolateral prefrontal cortex*, suggesting a need to suppress the L1 and greater attentional demands during L2 processing. Finally, the threshold for significant activation in the *superior temporal gyrus* was barely met, which the authors argued was due to impoverished semantic representations in the less proficient language, leading to diminished semantic retrieval and conceptual processing. Sebastian et al. (2011) draw a parallel to neural systems such as motor learning to support their conclusion.

While clear in focus and illustrative in its use of ALE methods, several critiques can be applied to this meta-analysis. Given Indefrey’s (2006) conclusions, it is noteworthy that five of the seven experiments in the analysis of greatest interest to the authors—L2 processing in the low proficiency group—comprise passive tasks, which may activate language areas poorly. The low proficiency group also contained a smaller number of studies, which may have contributed to the smaller and more distributed clusters. Finally, the selection of L2 proficiency as the only factor by which to compare studies may not have been optimal. Researchers (e.g., Abutalebi & Green, 2008) have shown that AoA plays a significant role in the organization of L1 and L2 processing, yet no interpretation of other relevant factors was attempted in the discussion. Finally, the number of foci and participants that each study contributed to the analysis was not reported. This is relevant because ALE meta-analyses with small data sets may be greatly skewed towards the findings of the two mostly highly powered studies (Eickhoff et al., 2016).

<u>Luk, Green, Abutalebi, & Grady (2012)</u> conducted an ALE meta-analysis comprised exclusively of bilingual neuroimaging studies that investigate language switching. The aim of this analysis was to empirically test the claims of Abutalebi and Green’s (2008) domain-general bilingual language control network. The authors selected studies with a variety of language switching tasks. Contrary to Sebastian et al.’s (2011) approach, they stipulated the use of a high-level baseline control condition designed to filter out non-linguistic processing that could be generated by the experimental condition. The final data set comprised ten experiments from ten studies, on which the authors performed just one analysis. The ALE calculations were corrected for multiple comparisons using the FDR algorithm with a somewhat more stringent threshold of *p* < 0.01 and a minimum cluster size of 100 mm^3^.

The analysis revealed clusters located primarily in the frontal regions of the left hemisphere. However, two regions named in Abutalebi & Green’s (2008) model—the *anterior cingulate cortex* and *bilateral supramarginal gryi*—were absent from their findings. The authors speculate that these regions may have escaped detection due to their decision to include only studies with matched control conditions. This illustrates the potential drawback of a high-level baseline: relevant data may be filtered out when fMRI activation in the original study fails to pass a significance threshold relative the control condition.

Luk et al. (2012) report the number of participants and foci per study and which studies contributed to each cluster. The authors illustrate an appropriate use of aggregating across tasks when the localization of a specific linguistic process or domain is not the intended goal. Here, greater variation in the purely linguistic elements of the data was advantageous, because the meta-analysis sought to isolate the language control component apart from linguistic processes per se. When there is little continuity between the linguistic components of tasks, it is less likely that foci will aggregate across studies and pass the significance threshold. Overall, Luk et al. (2012) confirms the effectiveness of a quantitative meta-analysis for testing theoretical models.

<u>Liu & Cao (2016)</u> investigated AoA as the primary factor of interest in their ALE meta-analysis. In addition, they questioned whether the differences that they identified in L1 and L2 networks might stem from dissimilarities between the written systems of each language, or from the effect of L2 learning on L1 processing. This metaanalysis is unique in considering linguistic differences between languages in terms of their orthographic depth, which refers to the processing levels required to access the phonological information represented by written symbols. Excluded studies, in addition to ROI studies and studies without coordinates or those that measured non-linguistic or language switching tasks, comprised studies with bilinguals outside of a stipulated age range (18-50). The authors gathered a data set of 57 studies with 102 experiments, using a minimum of 13 studies per analysis— the largest minimum to date—although the number of experiments taken from each study is not reported. ALE calculations were corrected for multiple comparisons using the FDR algorithm, a significance threshold of *p* < 0.05, and a minimum cluster volume of 150 mm^3^. Only clusters reported in at least two or three studies (when the total studies were < or > 15, respectively) were included in analysis (cf. Eickhoff et al., 2016).

The first analysis carried out by Liu & Cao (2016) addressed potential differences in the **L2 network by AoA**, when controlling for other participant characteristics. The authors only included foci from the L2>L1 contrast in contributing studies, excluding data from contrasts that utilized a baseline control. Therefore it is unsurprising that this meta-analysis generated smaller clusters than Sebastian et al. (2011). The results support previous studies that suggest the L2 network in late bilinguals recruits more neural resources, notably in frontal regions of the left hemisphere. Yet in this analysis, the *inferior frontal gyrus* unexpectedly yielded the smallest clusters in that area.

**Both groups** in separate analyses generated widely overlapping areas of elevated fMRI activation in the left hemisphere, specifically in frontal regions and the *insula*, although the coordinates in each language network represented neighboring rather than identical locations. A subtraction analysis between the two groups revealed elevated activation only in the *left superior frontal gyrus* for **late bilinguals**. The analysis for this group was the more highly powered of the two, with over twice as many subjects. Overall, the authors note the absence of clusters in temporal regions, similar to Indefrey (2006). Otherwise, the outer cortical regions that were identified closely resemble Luk et al.’s (2012) language control network. From Liu & Cao (2016) onward, consistent activation of inner cortical regions like the insula is found, perhaps reflecting a shift in task design within the literature.

Next the authors investigated the **L1 network by AoA**, comparing the L1>baseline contrast in early and late bilinguals. The number of studies with high and low proficiency level participants were balanced within each group. The authors do not specify why they chose the contrast with baseline controls for this analysis. Significant results are observed less often in the L1>L2 contrast, so potentially only this data type allowed for a sufficiently large data set to perform a fully-powered subtraction analysis. **Both groups** in separate analyses reveal clusters that are remarkably similar, and the subtraction analysis between them shows significantly elevated fMRI activation only in the left *fusiform gyrus* for **early bilinguals**.

Considering Sebastian et al.’s (2011) analysis of the same data type, it appears that AoA may be a less relevant factor than proficiency level for discerning an L1 network. The absence of clusters in the left *inferior frontal gyrus* is notable, but expected, as the area is implicated to a greater degree in L2 processing. The authors cite behavioral studies that allege early AoA might complicate the establishment of an independent L1 lexical system. However, the results contradict other behavioral studies that claim the L1 network is more profoundly affected in late bilinguals. The authors attribute this discrepancy to the heterogeneity in participant characteristics.

Finally, **orthographic depth** was investigated for its effect on the **L2 network**. Participant groups were carefully balanced by language type, AoA, and proficiency level. The **shallower L2** engaged the right hemisphere, largely in temporal regions, whereas the **deeper L2** generated results that resemble the AoA analyses, with clusters found largely in frontal regions of the left hemisphere. The authors concluded that different networks may form to process the same language based on its relation to the bilingual’s L1. They propose that orthographically shallower L2s may rely upon bilateral regions involved in phonological assembly, whereas deeper L2s may require greater involvement of left hemisphere regions subserving lexical integration. However, substantially more data was again available for late bilinguals.

While creative and careful in their approach and ambitious in scope, Liu & Cao (2016) may not have fully achieved their intended goals for several reasons. Firstly, a focus on orthographic depth as the most relevant linguistic difference between L1 and L2 presupposes that reading mediates language learning. The authors assume that additional processing effects are elicited by the language that is orthographically deeper, *even when the task lacks written stimuli items*. While many bilinguals may have learned their L2 in a formal classroom environment or may utilize their L2 for professional purposes that require reading and writing, this is unlikely to be representative of all bilinguals. In fact, AoA may represent a confound in this line of reasoning, as younger learners often learn to speak before learning to read, and heritage speakers may never become literate in their L1.

A second concern relates to the method of study selection: while rigorous in principle, numerous factors were in fact approximated by the authors. Studies were divided into three sets tailored to the goals of each analysis. *Set A* included the L2>L1 contrast from 40 studies to investigate the effect of AoA on the L2 network. Data from early and late bilinguals were compared in participants who were highly proficient in their L2, in order to control for this factor. The average participant age, the orthographic depth between languages, and the distribution of task types were estimated by the authors to be roughly equivalent across groups. Nonetheless, studies with late bilinguals (27 studies) were over-represented compared to those with early bilinguals (13 studies). *Set B* included the L1>baseline condition from 30 studies, this time evenly distributed between early and late bilinguals. The groups were again matched approximately by participant age, orthographic depth between languages, and range of task types. This set was created to investigate if AoA would affect the L1 network, so the number of L1 language types were matched across groups. *Set C* included the L2>baseline contrast from 50 studies to assess if the orthographic depth of L2 relative L1 would affect L2 processing. The two groups possessed an L2 that was either orthographically deeper or shallower than the L1 (25 studies each). The number of L2 languages represented, the average participant age, AoA, proficiency level, and range of task types were approximated across groups.

Although this meta-analysis is exemplary in its attempt to control for a wide range of factors known to affect language processing, ultimately the paper may provide a false sense of certainty in that a very large number of factors are nonetheless estimated or approximated, compounded by the fact that studies are heterogenous in their labeling practices for participant information, which means that different factors may intersect in unforeseen ways which cannot be reported. A Chi square test was employed to estimate the degree of similarity between tasks in each subset, but the test still required a subjective labeling process to compile the tasks into articulatory, phonological, orthographic, semantic, or “sentential” categories. Thus, characteristics assumed to be “matched” across groups may in fact be over-represented in one task. The strength of a meta-analysis lies in aggregating actual data to create a larger sample than obtained in individual studies, such that data need not be approximated. In this sense, the meta-analysis may inadvertently resemble Luk et al.’s (2012) approach: the heterogeneity in each subset may average out information relevant to each intended analysis, revealing some shared linguistic factor, but one that is difficult to define. The small number of clusters obtained and the unexpected findings noted by the authors support this interpretation. Nonetheless, Liu & Cao (2016) provide interesting exploratory findings that foreshadow the type of meta-analysis that could be conducted when a richer data set becomes available.

<u>Cargnelutti, Tomasino, & Fabbro (2019)</u> also prioritize AoA in their ALE meta-analysis, however these authors focus on the lower bound for the onset of a critical period for language acquisition. They compare early bilinguals (up to 6 years) to very early bilinguals (up to 3 years), a distinction motivated by the timeline for development of linguistic memory systems. A secondary aim was to control for linguistic skill level by investigating AoA in the group of highly proficient bilinguals. For their first analysis, the authors collected the same size data set as Liu & Cao (2016), but allowed more contrasts—and therefore more data—per study. This comprised a total of 57 studies with 17 experiments (8 studies) for very early bilinguals, 74 experiments (12 studies) for early bilinguals, and 174 experiments (37 studies) for late bilinguals. ROI studies and studies without coordinates were excluded, in addition to those with participants who were bimodal bilinguals or with AoA that did not correspond to the authors’ classification scheme. Studies that required a very low level of linguistic processing, such as passive viewing of letters, were not analyzed. The primary analysis allowed all data types that included any linguistic data, even when the contrast was between tasks, rather than between languages. In the second analysis, the authors utilized the L1 or L2>baseline contrast and rejected complex tasks that spanned linguistic and non-linguistic competencies, such as translation and language switching. However, exceptions to these criteria were noted. ALE calculations were corrected at the cluster level for multiple comparisons using a significance threshold of *p* < 0.05 and a minimum cluster volume of 200 mm^3^ for main effects and a reduced cluster volume of 80-120 mm^3^ for other comparisons. The significance threshold was reduced to *p* < 0.001 (uncorrected) for contrast analyses.

In their first meta-analysis, the authors provide a general overview of language processing by aggregating all available data types that involved either L1 or L2. This **“combined” language network** is compared between the very early, early, and late bilingual groups. Substantial overlap was found for the location of increased fMRI activation in the two older groups, although the extent of this activation differs strikingly across all three groups, increasing in scope across both hemispheres as AoA rises. **Very early bilinguals** generated just three clusters in the left *middle temporal gyrus* and bilateral *cerebella*. The cluster in the left *cerebellum* was the only cluster unique to this group. However, upon closer inspection of the studies in this data set, only five of eight studies compare data between languages; the others compare between tasks, between monolinguals and bilinguals, or represent language switching. This presents the question of how meaningful the results may be compared to those of bilinguals with a later AoA, for whom more stringent data exclusion criteria were adopted. Additionally, given the very small number of clusters obtained for very early bilinguals when combining across languages, it is questionable whether unique results would be obtained for L1 and L2 independently, even in a larger data set.

The **late and early groups** of bilinguals generated very similar networks in the left hemisphere, with clusters located throughout traditional language areas. Results specific to each group of bilinguals predominately represent neighboring or adjacent regions, or a change in laterality. A minor shift in coordinates could be attributed to individual participant differences in the underlying data set. This is less likely for changes in laterality.

Although clusters span similar regions in the left hemisphere, the greater extent of activation found in frontal areas for **late bilinguals** is notable.

This group also reveals more extensive activation in the right hemisphere. Indefrey (2006) draws a connection between right hemisphere activation and certain task types. A confound therefore may be created by unbalanced distribution of tasks across participant groups.

Overall, the results suggest that AOA after the age of six has little effect on which regions comprise a combined language network. This conclusion must remain tentative, however, given the heterogenous and minimal data available for very early bilinguals.

In their next analysis, the authors investigated the L1 and L2 networks separately, utilizing L1 or L2>baseline contrasts. The majority of studies with very early bilinguals lacked feasible data for this contrast, which is likely why the group was excluded from further analyses. However, the very early and early AoA data sets are combined in these later analyses. The combined data set is still roughly half the size of the late AoA data set.

In the **L1 network**, clusters were again obtained for **both groups** throughout the left hemisphere. These regions largely fell within the traditional language areas; however, larger clusters were observed in frontal regions for **late bilinguals**, and in posterior regions for **early bilinguals**.

In the **L2 network**, the same frontal and posterior regions were activated in **both groups** in the left hemisphere; however, this time more extensive activation was found for **late bilinguals** in posterior regions. Unique activation for **early bilinguals** was found only the left *precentral gyrus*. With and between group comparisons for both language and participant groups revealed minimal results, centered around the *inferior frontal gyrus*.

Thus far, the meta-analysis conducted by Liu & Cao (2016) is the closest in design and aim to Cargnelutti et al. (2019). Many clusters identified here lie adjacent to areas activated in Liu & Cao (2016), but with more extensive results than those obtained in the earlier meta-analysis. In the **L1 network**, the only regions common to both studies include the left *precentral gyrus*, despite the use of the same data type by both authors. In the **L2 network,** the earlier authors used the more stringent L2>L1 contrast, which might explain why fewer regions overall were identified. Nonetheless, this contrast yielded a greater number of regions common to both studies: the left *inferior frontal gyrus* and *precentral gyrus* for **early bilinguals**, and the left *inferior frontal gyrus, medial frontal gyrus*, and *insula* for **late bilinguals**. In accordance with previous findings, late bilinguals elicited clusters that were larger in size, more varied in their distribution, and located bilaterally for **both L1 and L2 networks**.

Next the authors investigated AoA among just the subset of highly proficient bilinguals in attempt to control for the effect of proficiency level. In this analysis, the data sets for early and late bilinguals were balanced with 16 and 17 studies in each group. In the **L1 network**, clusters were found exclusively in the left hemisphere, again primarily in frontal regions. **Early bilinguals** also activated the *middle temporal gyrus*. In the **L2 network**, only the *inferior frontal gyrus* was found for **early bilinguals**, whereas **late bilinguals** produced a variety of clusters in both hemispheres. A finding for this group that was unique among all of the other analyses in the paper was activation of the *caudate nucleus*, which is a part of the language control network. Within and between group analyses revealed results only for conjunction analyses: the left *inferior frontal gyrus* for the **L2 network** of **both groups**, and the left *inferior frontal gyrus* for **L1 and L2 in proficient late bilinguals**.

Sebastian et al. (2011) provides the most relevant comparison for this analysis. The earlier study uses the same data type, but considers proficiency level independent of AoA. This time, the two meta-analyses come to strikingly different conclusions: not one region is activated in common, and the subtraction analyses do not replicate. The **L1 network** is more extensive in Sebastian et al. (2011), while this is true for the **L2 network** in Cargnelutti et al. (2019). For **both networks**, the earlier study identifies clusters outside of core language areas, and the later study locates clusters within the classical language network. These findings may show AoA to be a more relevant factor than proficiency level or result from the use of a larger data set in Cargnelutti et al. (2019).

The overall impression produced by Cargnellutti et al.’s (2019) meta-analyses is that differences between early and late bilinguals lie primarily in the scope of activation, rather than in the core regions associated within L1 and L2 networks. Right hemisphere clusters yield some of the most substantial differences between groups, yet these results are not replicated in earlier meta-analyses. The study did uphold previous findings that the L1 network in early and late bilinguals differ substantially, and that while activation found in the L2 network for late bilinguals is wider and more varied, it largely encompasses the same cortical regions as the L1 network. The larger data set analyzed by these authors should provide more power for conjunction and subtraction analyses. Therefore, the lack of findings outside of the left *inferior frontal gyrus* indicates that differences between the language networks may indeed stem from the extent and manner of activation for each language (cf. Hernandez, Claussenius-Kalman, Ronderos, Castilla-Earls, Sun, Weiss, & Young, 2019) rather than from the individual areas that comprise the networks. However, the fact that exclusion criteria do not appear to be held constant across groups is a serious concern, particularly when the heterogenous data from very early bilinguals is merged with an already small sample of early bilinguals. Considering the variety of data types combined, there is the risk, similar to Liu & Cao’s (2016) analysis, that this heterogeneity may prevent isolation of truly linguistic results. Nonetheless, Cargnellutti et al. (2019) show that an ALE meta-analysis, perhaps in conjunction with larger data sets, do appear to isolate more relevant brain regions for L1 and L2 processing than Indefrey’s (2006) more subjective method.

<u>Sulpizio, Del Maschio, Fedeli, & Abutalebi (2020)</u> return to linguistically-defined task distinctions as the organizing principle for their ALE meta-analysis. Unlike Indefrey (2006), the authors do not consider which processing levels may be required by specific task designs and do not differentiate production and comprehension tasks. Instead, linguistic domains are defined broadly in three categories: *lexico-semantics* describes tasks that involve the association of meaning to a linguistic structure; *grammar* describes tasks that involve the recognition and production of grammatical structures; and *phonology* combines tasks that involve the recognition and production of sound patterns. *Language switching* studies are analyzed as a fourth category. The two domains that comprise the largest data sets—lexico-semantics and language switching—are further broken down by AoA in a secondary analysis. With 52 studies, this meta-analysis is comparable in scope to other recent publications. However, even with quite broadly defined domains, a number of analyses make use of as little as five studies. The number of experiments and foci taken from each study are not specified. The number of foci and contrasts taken from each paper is reported instead. Excluded studies did not publish coordinates reported in Montreal Neurological Institute (MNI; Collins et al., 1998) stereotaxic space, did not target linguistic tasks, analyzed ROIs, or presented results from a participant’s third or subsequent language. The authors report that only the L2>L1 or L1>L2 contrasts comprise the data set for the domain analyses, although exceptions to this criterion can be found. All analyses were conducted with an uncorrected threshold of *p* < .001 and a minimum cluster size of 150mm^3^.

The analysis of the **lexico-semantic domain** produced the largest network of all categories, with clusters located throughout frontal and posterior regions in both hemispheres. This finding is unsurprising, considering the heterogenous nature of the data set, which encompassed a huge range of task types and underlying processes. For example, all of the tasks analyzed in Indefrey (2006) except sentence level listening and reading fall into Sulpizio et al.’s (2020) lexico-semantic domain.

In general, the results of the analysis lie in stark contrast to previous studies: the **L1 network** was found to be more extensive and bilateral in composition than the L2 network. The regions identified included a wide range of subcortical areas located outside of the language control network, such as the *thalamus* and *amygdala*. This was also true for the **L2 network**, which included the *cerebellum* and bilateral *insulae* and *pallidum*. However, two clusters from Indefrey (2006) are replicated: BA 21 in the L1 network and BA 44/45 in the L2 network. The overlap between **both networks** in the *left inferior frontal gyrus* is also shared with previous meta-analyses.

Similar trends are observed in the **grammatical domain**. Once again, only the *inferior frontal gyrus* was common to **both networks**. Unique findings for this domain include the *bilateral insulae*, which, contrary to the lexico-semantic analysis, have now shifted to the **L1 network**, and the L2 network was almost entirely left-lateralized. The tasks in this domain are similar to Indefrey’s (2006) sentence level reading and listening category, which required a grammatical decision. The only cluster to replicate between analyses was the *inferior frontal gyrus*.

The **phonological domain** was the only category to produce more extensive activation in the **L2 network**, primarily in frontal regions with just one posterior cluster in the *superior parietal lobule*. The **L1 network** centered around the right *temporal gyrus*. For **both networks**, the only region identified was the left *inferior frontal gyrus*. While the previous two domains compared relatively balanced data sets, the L1 network here was derived from 5 studies, and the L2 from 10 studies. Previous meta-analyses have not investigated phonology as an independent category, which prevents a comparative evaluation of the results.

**Language switching** studies were analyzed relative a baseline condition. Clusters appeared in an extensive bilateral network with widespread activation throughout frontal and posterior regions. Bilateral clusters were found only in the *inferior frontal gyrus* and the *precuneus*. The left hemisphere comprised largely posterior regions, and the right hemisphere generally exhibited more frontal regions; this appears to be a trend, with similar findings in two of the three other linguistic domains. Subcortical activation was found only in the left hemisphere. Two of the clusters reported in Luk et al. (2012) are replicated here in the left hemisphere, and just one cluster in the right hemisphere. Given the careful selection criteria adhered to in Luk et al. (2012), the divergent results obtained by Sulpizio et al. (2020) may represent activation in areas outside of the language control network.

A secondary analysis investigating the **effect of AoA** was undertaken for the lexico-semantic and language switching domains. Within the **lexico-semantic domain**, the **L1 network** of **early bilinguals** reproduced the widespread activation found when averaging across AoA, with the exception of the right *caudate* and the addition of the left *fusiform gyrus*. **Late bilinguals** generated three clusters: the bilateral *inferior frontal gyrus* and right *caudate*. In the **L2 network**, it was **early bilinguals** who revealed the smaller network in the left *inferior frontal gyrus*, *left insula*, and right *middle occipital gyrus*. **Late bilinguals** produced a wider range of clusters in frontal regions and interior cortical regions. Relative one another, these results appear logical.

Sebastian et al. (2011) found greater activation in the L1 network for highly proficient bilinguals, and this factor may play a role in explaining Sulpizio et al.’s (2020) findings. The authors state they included only studies with highly proficiency bilinguals, yet they acknowledged difficulties in establishing objective criteria by which to evaluate participant characteristics. Their findings show an extensive and bilateral network recruited for L1 processing, which contradicts most previous findings. This could also lend support to theories that the L2 affects the semantic representations of the L1 (Titone, Libben, Mercier, Whitford, & Pivneva, 2011).

**Language switching** tasks yielded substantial bilateral activation for both participant groups. The clusters found when averaging across AoA were distributed evenly between the groups. fMRI activation spanned frontal and posterior regions, with the occasional new cluster appearing in frontal, posterior, and subcortical regions. In the left hemisphere, **both groups** produced similar clusters in the *inferior frontal gyrus, precentral gyrus*, and *cerebellum*, and in the *inferior frontal gyrus* and *cuneus* in the right hemisphere. Bilateral results were found in frontal regions for **early bilinguals**, and in frontal and posterior areas for **late bilinguals**.

Neither group fully replicated Luk et al.’s (2012) language control network. **Early bilinguals** recruited a larger number of control regions. The literature predicts **late bilinguals** may require greater neural resources to suppress their L1 (cf. Abutalebi & Green, 2007), yet this group activated just one region from the earlier study: the *supplementary motor area*. A second cluster was found in the *anterior cingulate gyrus*. While absent from Luk et al. (2012), this region is part of the language control network described by Abutalebi and Green (2008).

Isolated results from Sulpizio et al. (2020) support previous research findings, but overall the study generates more contradictions of accepted beliefs in the bilingual neuroimaging literature. It may be, as the authors claim, that they have produced the first truly comprehensive bilingual ALE meta-analysis and therefore their data has greater validity. However, the authors themselves express difficulty in interpreting some of their controversial findings, and nearly fifteen years later, the number of studies per analysis or linguistic domain remains in some analyses just as limited as in Indefrey (2006). Most troublingly, doubts as to the rigor of the study selection criteria persist: a close reading of studies that comprise the lexico-semantic domain found that no less than 7 of 18 studies, or nearly 40% of the total data set, appear to be misclassified in respect to the data, participant, or task type. If categorization errors of this nature exist throughout their corpus, it would strongly undermine the reliability of the findings. For this reason, we refrain from drawing strong conclusions based on this meta-analysis.

This short review of bilingual meta-analyses has shown that, despite the explosive growth in interest in bilingual neuroimaging, considerable work is still required to generate an appropriate data set for meta-analyses, and as a consequence, to ultimately establish certain facts about L1 and L2 processing. Even allowing for very broadly defined categories, the available data is insufficient to make fine-grained comparisons or test hypotheses with multiple factors. A second important conclusion concerns the nature of conducting a meta-analysis in a field with an admittedly complex literature. The task of accurate categorization cannot be understated; many bilingual neuroimaging studies do not present their underlying assumptions, criteria, or design features clearly, if at all. As studies become more ambitious in their aim to address multiple factors, the actual details of their method for study selection and the technical specifications for each individual contrast within the paper become harder to discern. A final responsibility lies with researchers to publish data from all contrasts, even if they do not directly pertain to the primary aim of the study, to adopt clear standards of reporting, and to acknowledge study limitations.

Nonetheless, all of the meta-analyses that were reviewed presented a well-researched and largely convincing justification of their results. Yet in assessing the overall scope of these meta-analyses, we can conclude that there is both the lack of a common interpretation that would explain differences in L1 and L2 processing, and a lack of evidence that there is indeed no difference between L1 and L2 processing. It is possible to have numerous different accounts that are all well-argued by drawing upon the rich, but sometimes conflicting results that can be found from individual studies in the monolingual and bilingual neuroimaging literature. Somewhat counter-intuitively, these meta-analyses make little reference to one another. Therefore, in our meta-analysis, rather than justifying our results with findings from individual neuroimaging studies, we will directly compare our results to those of earlier bilingual neuroimaging meta-analyses, with an account of what decisions in our experimental design and analysis parameters may have led to findings that are consistent with or divergent from previous work. We will attempt to convey in a transparent fashion what is possible to achieve in a task-based meta-analysis of bilingual neuroimaging studies at this point in time, and what shortcomings may be discerned in the method and data set available. Furthermore, activation in key language regions such as the left *inferior frontal gyrus* is the most consistent finding across meta-analyses, yet each study makes a case for a unique set of regions dedicated to L2 processing. Therefore, our analysis will take a greater interest in where L1 and L2 networks appear to be disassociated and what evidence there maybe for linguistic processing outside of core language regions.

## 2. Method

### 2.1 Paper Selection

Our study selection process is documented in a PRISMA flowchart (Moher et al., 2009; Page et al., 20201; Figure 1). PRISMA, or Preferred Reporting Items for Systematic Reviews and Meta-Analyses, provides a standardized format for describing how data sources are identified and at what stage and why studies are discarded from the analysis. The initial stage of data collection identified the greatest number of potentially relevant studies within the Medline and Scholar online databases, utilizing two broad search terms (fMRI, bilingual) for the time period through the end of 2020. Additionally, the reference section of the relevant studies obtained through this process and from previous bilingual neuroimaging meta-analyses were consulted, but provided no new entries found outside of the original query.

**Figure 1.**
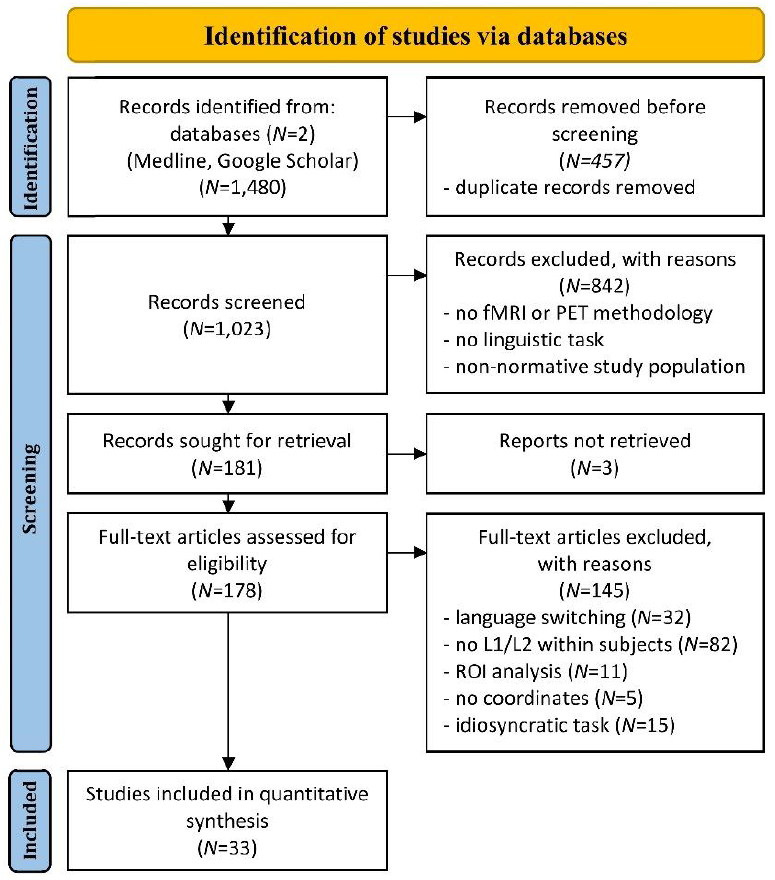
PRISMA flow chart of the paper selection process.

**Figure 2.**
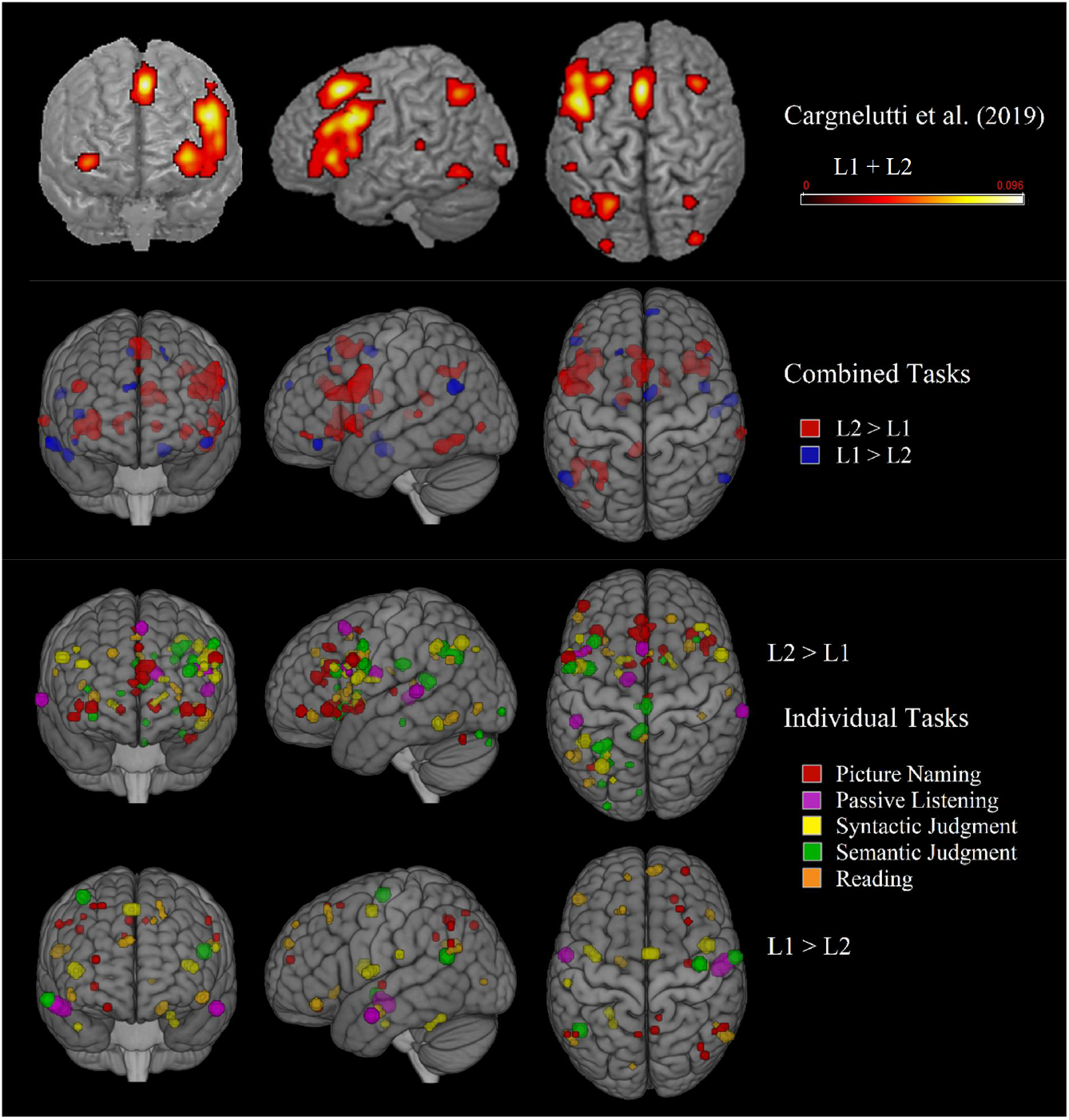
Combined and Aggregated Results Note: Results for the five individual tasks are show b) combined and c) aggregated. These results fall within the same regions identified in a) the combined L1+L2 network from a recent meta-analysis with no task distinction. The uppermost row of pictures is reproduced with the author’s permission from Cargnelutti et al. (2019).

**Figure 3.**
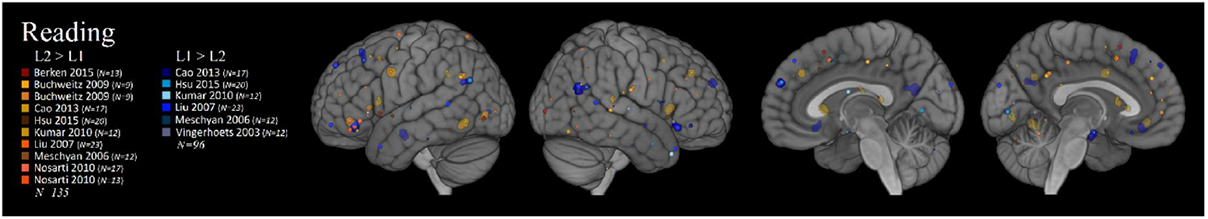
Reading Task: Individual and Cumulative Results Note: Foci from individual experiments are superimposed over fMRI activation clusters from the meta-analysis.

**Figure 4.**
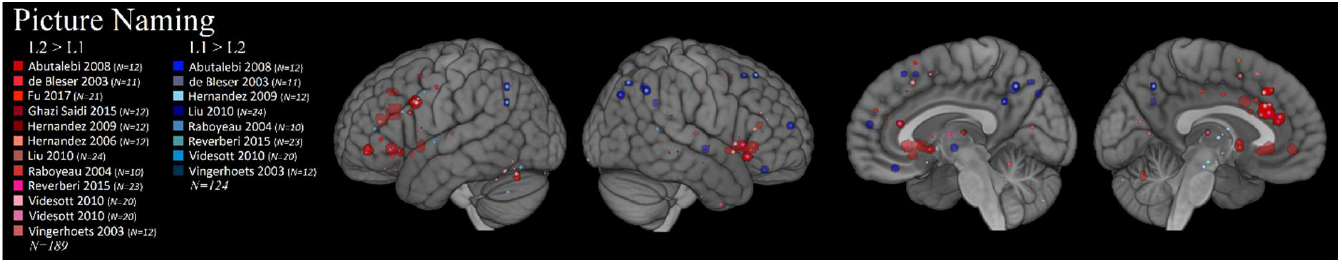
Picture Naming Task: Individual and Cumulative Results Note: Foci from individual experiments are superimposed over fMRI activation clusters from the meta-analysis.

**Figure 5.**
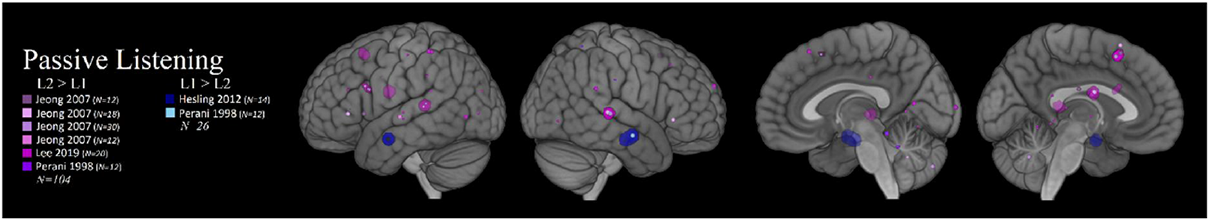
Passive Listening Task: Individual and Cumulative Results Note: Foci from individual experiments are superimposed over fMRI activation clusters from the meta-analysis.

**Figure 6.**
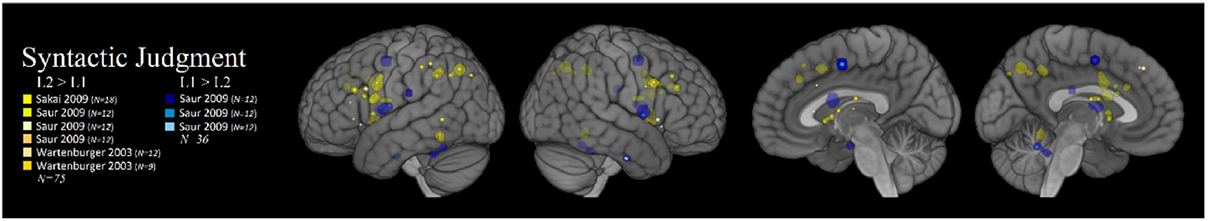
Syntactic Grammaticality Judgment: Individual and Cumulative Results Note: Foci from individual experiments are superimposed over fMRI activation clusters from the meta-analysis.

**Figure 7.**
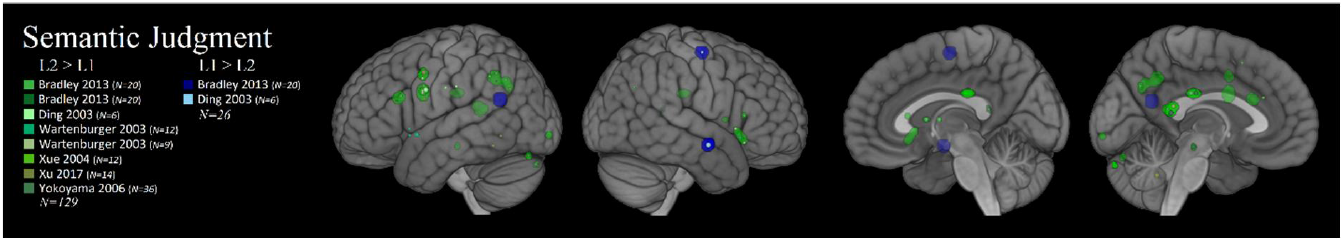
Semantic Similarity Judgment: Individual and Cumulative Results Note: Foci from individual experiments are superimposed over fMRI activation clusters from the meta-analysis.

**Figure 8.**
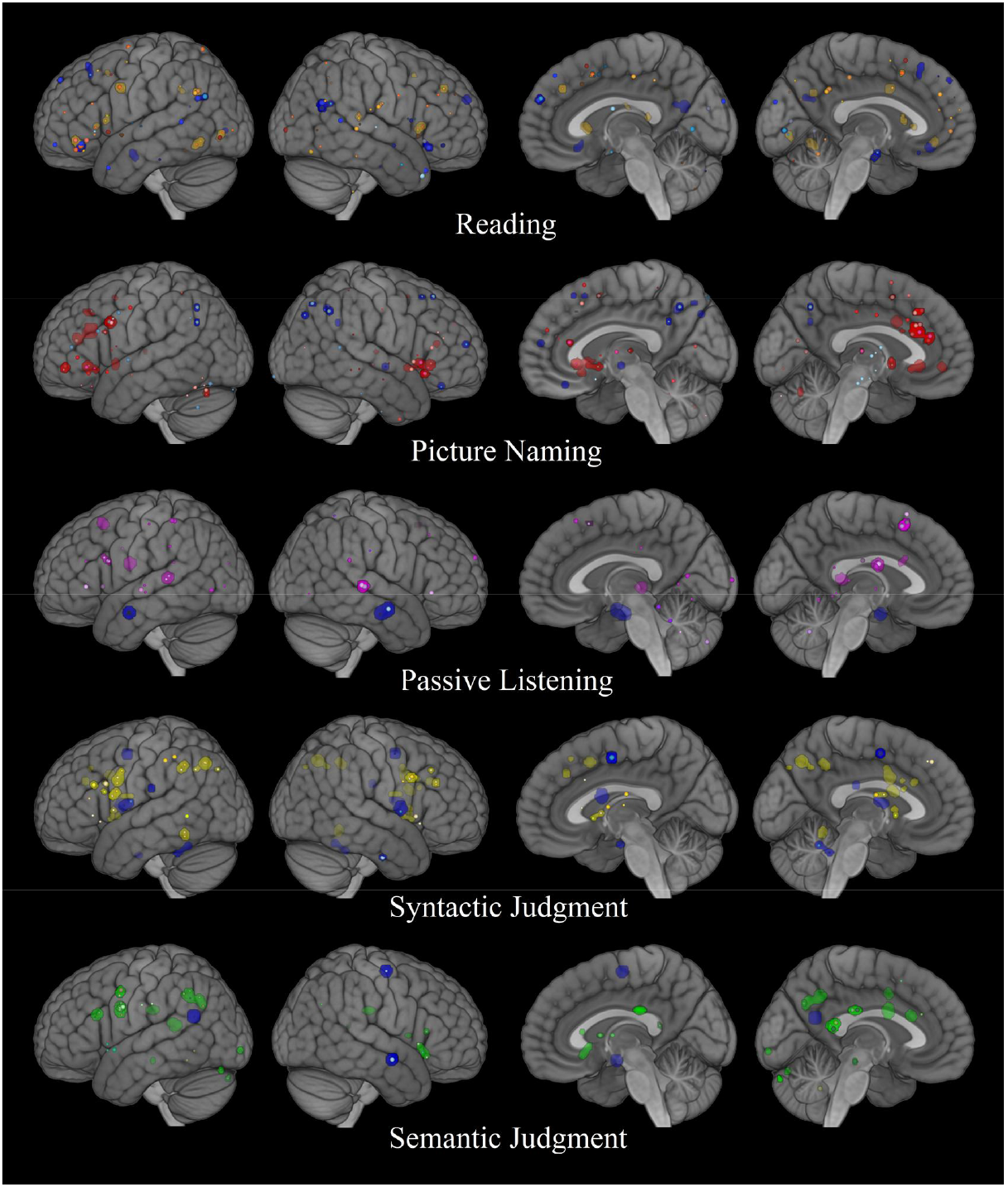
Five Tasks: Overview Note: Blue represents the L1>L2 contrast. All other colors represent the L2>L1 contrast in their respective tasks

The initial screening of records excluded all studies with a focus and methodology that would not provide data suitable for our research questions. This included studies with no fMRI or PET component, those without a linguistic task or with additional confounds that did not allow a for the clear designation of a linguistic task (e.g., Li et al., 2013), research investigating non-verbal modalities of language, or carried out among a non-normative study population (such as those with aphasia, dyslexia, etc.).

We next evaluated the eligibility of the remaining full-text articles according to several criteria. Studies that could be excluded according to more than one criterion were evaluated in the order presented here. Studies with language switching paradigms were discarded, followed by those that did not report a contrast between L1 and L2. This eliminated all studies that provided data only for contrasts of L1 or L2 relative a baseline condition, studies that contrasted bilinguals with monolinguals, and analyses that were conducted between subjects rather than within subjects or that combined data collected in both languages. The remaining studies possessed one task per contrast with the task performed separately in each language and with a main effect of language.

Next, studies that only reported findings from ROIs were removed from consideration. The use of ROIs reflects an a priori decision on the part of researchers about where results are expected to be found and creates a more lenient statistical threshold for fMRI activations to be considered significant; inclusion of these studies would have allowed data that represented different assumptions into the meta-analysis, giving greater a weight to ROI studies that may not be warranted. Studies that reported whole brain L1 versus L2 contrasts in addition to ROI analyses were retained, but only the former data points were entered into the meta-analysis. Finally, studies that did not publish a full set of stereotactic coordinates for their findings were necessarily excluded from the analysis.

The remaining 48 studies were assessed for the relative similarity of their task designs, of which 33 were retained in five categories: *reading (10), picture naming (11), passive listening (5), semantic similarity judgment (6)*, and *syntactic grammaticality judgment (3)*. Several studies contributed more than one experiment within the same or different task categories (see Table 1). No task considered less than eight experiments: *reading (13), picture naming (12), passive listening (10), semantic similarity judgment (9)*, and *syntactic grammaticality judgment (8)*. It is of particular note that the last five years have yielded only four new studies suitable for inclusion into a meta-analysis with these criteria.

**Table 1.** Indefrey (2006) Results

| **Semantic Decision** |  |  |
| --- | --- | --- |
| L1>L2 |  |  |
| LR | Anatomical Region | BA |
| L | Anterior Middle Temporal Gyrus | 21 |
| L2>L1 |  |  |
| LR | Anatomical Region | BA |
| L | Posterior Middle Frontal Gyrus | 9 |
| L | Posterior Inferior Frontal Gyrus | 45 |
| L | Posterior Inferior Parietal Lobule | 39 |
| L | Anterior Cingulate |  |
| **Word Generation** |  |  |
| L2>L1 |  |  |
| LR | Anatomical Region | BA |
| L | Posterior Inferior Frontal Gyrus | 47 |
| R | Posterior Inferior Frontal Gyrus | 47 |
| **Picture Naming** |  |  |
| L2>L1 |  |  |
| LR | Anatomical Region | BA |
| L | Posterior Inferior Frontal Gyrus | 44/47 |
| **Sentence Listening/Reading** |  |  |
| L2>L1 |  |  |
| LR | Anatomical Region | BA |
| L | Posterior Middle Frontal Gyrus | 9 |
| L | Posterior Inferior Frontal Gyrus | 44/47 |
|  | Supplementary Motor Area |  |

*Note: Bilateral results are highlighted*

**Table 2.** Sebastian et al. (2011) Results

| **High Proficiency**<br>Both L1 and L2 (>baseline) |  |  | **Low Proficiency**<br>Both L1 and L2 (>baseline) |  |  |
| --- | --- | --- | --- | --- | --- |
| LR | Anatomical Region | BA | LR | Anatomical Region | BA |
| L | Inferior Frontal Gyrus | 47 | L | *Inferior Frontal Gyrus* | 45 |
| L | *Inferior Frontal Gyrus* | 45 | L | *Inferior Frontal Gyrus* | 44 |
| L | *Inferior Frontal Gyrus* | 44 | L | *Superior Temporal Gyrus* | 22 |
| L | Superior Frontal Gyrus | 8 | L | Angular Gyrus | 39 |
| L | Middle Temporal Gyrus | 21 | L | *Fusiform Gyrus* | 37 |
| L | *Superior Temporal Gyrus* | 22 | L | Occipital Gyrus | 18 |
| L | *Fusiform Gyrus* | 37 |  |  |  |
| L | Cerebellum |  |  |  |  |
|  |  |  | R | Occipital Gyrus | 18 |
|  |  |  | R | Cerebellum |  |
| Only L1 (L1>baseline) |  |  | Only L1 (L1>baseline) |  |  |
| LR | Anatomical Region | BA | LR | Anatomical Region | BA |
| L | Precentral Gyrus | 4 | L | Middle Temporal Gyrus | 21 |
| L | Precuneus | 7 | L | Supramarginal Gyrus | 40 |
| L | Occipital Gyrus | 19 | L | Occipital Gyrus | 19 |
| L | Occipital Gyrus | 18 |  |  |  |
| L | Cuneus | 17 |  |  |  |
| R | Cerebellum |  | R | Middle Temporal Gyrus | 21 |
| Only L2 (L2>baseline) |  |  | Only L2 (L2>baseline) |  |  |
| LR | Anatomical Region | BA | LR | Anatomical Region | BA |
| L | *Middle Frontal Gyrus* | 46 | L | Inferior Frontal Gyrus | 47 |
|  |  |  | L | *Middle Frontal Gyrus* | 46 |
|  |  |  | L | Middle Frontal Gyrus | 9 |
|  |  |  | L | Paracingulate Gyrus | 32 |
|  |  |  | L | Cingulate Gyrus | 24 |
|  |  |  | L | Precentral Gyrus | 4 |
|  |  |  | L | Precuneus | 7 |
| R | Temporal Pole |  | R | Inferior Frontal Gyrus | 45 |
|  |  |  | R | Clastrum |  |
|  |  |  | R | Precuneus | 7 |

*Note: Bilateral clusters are highlighted. Between-analysis clusters are italicized. The data type used in each analysis is indicated in parentheses.*

**Table 3.** Luk et al. (2012)

| Language Switching (>baseline) |  |  |
| --- | --- | --- |
| LR | Anatomical Region | BA |
| L | Middle Frontal Gyrus | 9 |
| L | Middle Frontal Gyrus | 46 |
| L | Inferior Frontal gyrus | 47 |
| L | Inferior Frontal Gyrus | 44 |
| L | Middle Temporal Gyrus | 37 |
| L | Caudate |  |
|  | Midline Pre-SMA | 6 |
| R | Precentral Gyrus | 6 |
| R | Superior Temporal Gyrus | 22 |
| R | Caudate |  |

*Note: Bilateral results are highlighted. The data type is indicated in parentheses.*

**Table 4.** Liu & Cao (2016) Results

| **Only Early (L2>L1)** |  |  | **Only Late (L2>L1)** |  |  |
| --- | --- | --- | --- | --- | --- |
| LR | Anatomical Region | BA | LR | Anatomical Region | BA |
| L | *Middle Frontal Gyrus* | 10 | L | *Middle Frontal Gyrus* | 9 |
| L | *Inferior Frontal Gyrus* | 9 | L | *Medial Frontal Gyrus* | 8 |
| L | *Precentral Gyrus* | 6 | L | *Superior Frontal Gyrus* | 6 |
| L | *Insula* | 13 | L | *Insula* | 47 |
|  |  |  | L | *Inferior Frontal Gyrus* | 46 |
|  |  |  | L | *Precentral Gyrus* | 44 |
| **Early vs. Late (L2>L1)** |  |  | **Late vs. Early (L2>L1)** |  |  |
| LR | Anatomical Region | BA | LR | Anatomical Region | BA |
|  |  |  | L | *Superior Frontal Gyrus* | 8 |
|  |  |  | L | *Superior Frontal Gyrus* | 6 |
| **Only Early (L1>baseline)** |  |  | **Only Late (L1>baseline)** |  |  |
| LR | Anatomical Region | BA | LR | Anatomical Region | BA |
| L | *Medial Frontal Gyrus* | 6 | L | *Superior Frontal Gyrus* | 6 |
| L | *Precentral Gyri* | 6 | L | *Medial Frontal Gyrus* | 6 |
| L | *Precentral Gyri* | 4 | L | *Precentral Gyri* | 6 |
| L | *Superior Temporal Gyrus* | 22 | L | *Precentral Gyri* | 4 |
| L | *Fusiform Gyrus* | 37 | L | *Superior Temporal Gyrus* | 38 |
| L | *Superior Parietal Lobule* | 7 | L | *Superior Temporal Gyrus* | 22 |
|  |  |  | L | *Precuneus* | 7 |
| **Early>Late L1 (L1>baseline)** |  |  | **Late>Early L1 (L1>baseline)** |  |  |
| LR | Anatomical Region | BA | LR | Anatomical Region | BA |
| L | *Fusiform Gyrus* | 37 |  |  |  |
| **Shallower>Deeper L2 (L2>baseline)** |  |  | **Deeper>Shallower L2 (L2>baseline)** |  |  |
| LR | Anatomical Region | BA | LR | Anatomical Region | BA |
| L | *Middle Temporal Gyrus* | 21 | L | *Inferior Frontal Gyrus* | 47 |
|  |  |  | L | *Middle Frontal Gyrus* | 46 |
|  |  |  | L | *Inferior Frontal Gyrus* | 45 |
|  |  |  | L | *Medial Frontal Gyrus* | 8 |
|  |  |  | L | *Insula* | 13 |
| R | *Precentral Gyrus* | 6 |  |  |  |
| R | *Superior Temporal Gyrus* | 21 |  |  |  |

*Note: Between-analysis clusters are italicized. The data type is indicated in parentheses.*

**Table 5.**
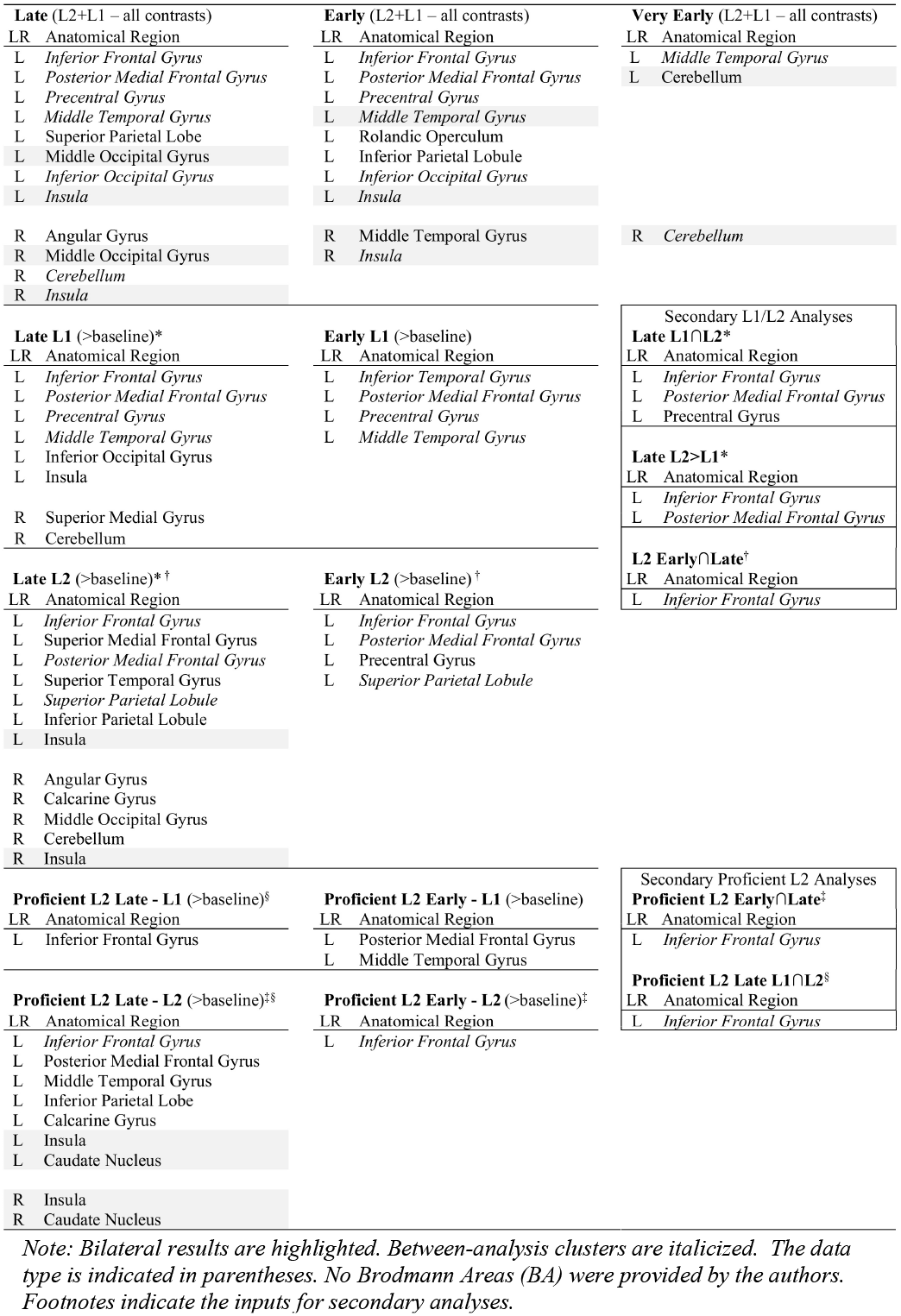
Cargnelutti et al. (2019) Results

**Table 6a.**
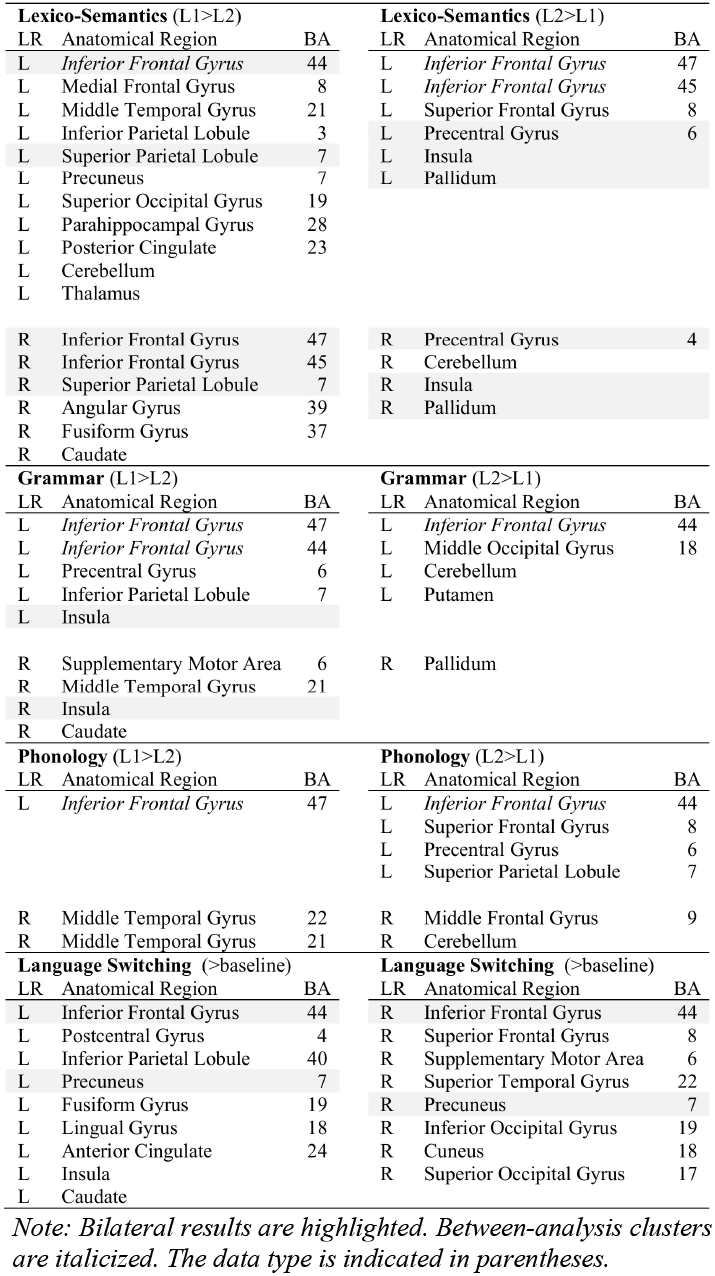
Sulpizio et al. (2020) Results

**Table 6b.** Sulpizio et al. (2020) Results

| Lexico-Semantics Early (L1>L2) |  |  | Lexico-Semantics Late (L1>L2) |  |  |
| --- | --- | --- | --- | --- | --- |
| LR | Anatomical Region | BA | LR | Anatomical Region | BA |
| L | *Inferior Frontal Gyrus* | 44 | L | *Inferior Frontal Gyrus* | 47 |
| L | Medial Frontal Gyrus | 8 |  |  |  |
| L | Middle Temporal Gyrus | 21 |  |  |  |
| L | Inferior Parietal Lobule | 3 |  |  |  |
| L | Precuneus | 31 |  |  |  |
| L | Precuneus | 7 |  |  |  |
| L | Fusiform Gyrus | 37 |  |  |  |
| L | Parahippocampal Gyrus | 34 |  |  |  |
| L | Parahippocampal Gyrus | 28 |  |  |  |
| L | Posterior Cingulate | 23 |  |  |  |
| L | Cerebellum |  |  |  |  |
| L | Thalamus |  |  |  |  |
| R | *Inferior Frontal Gyrus* | 47 | R | *Inferior Frontal Gyrus* | 38 |
| R | *Inferior Frontal Gyrus* | 45 | R | Caudate |  |
| R | Superior Parietal Lobule | 7 |  |  |  |
| R | Angular Gyrus | 39 |  |  |  |
| R | Fusiform Gyrus | 37 |  |  |  |
| Lexico-Semantics Early (L2>L1) |  |  | Lexico-Semantics Late (L2>L1) |  |  |
| LR | Anatomical Region | BA | LR | Anatomical Region | BA |
| L | *Inferior Frontal Gyrus* | 45 | L | *Inferior Frontal Gyrus* | 47 |
| L | *Insula* |  | L | *Inferior Frontal Gyrus* | 45 |
|  |  |  | L | Superior Frontal Gyrus | 8 |
| R | Middle Occipital Gyrus | 19 | L | Precentral Gyrus | 6 |
|  |  |  | L | *Insula* |  |
|  |  |  | L | *Pallidum* |  |
|  |  |  | R | Cerebellum |  |
|  |  |  | R | *Insula* |  |
|  |  |  | R | *Pallidum* |  |
| Language Switching Early (>baseline) |  |  | Language Switching Late (>baseline) |  |  |
| LR | Anatomical Region | BA | LR | Anatomical Region | BA |
| L | Middle Frontal Gyrus | 46 | L | *Inferior Frontal Gyrus* | 44 |
| L | *Inferior Frontal Gyrus* | 44 | L | *Precentral Gyrus* | 4 |
| L | *Precentral Gyrus* | 6 | L | Inferior Parietal Lobule | 40 |
| L | Fusiform Gyrus | 19 | L | Precuneus | 7 |
| L | Middle Occipital Gyrus | 18 | L | Angular Gyrus | 39 |
| L | *Cerebellum* |  | L | Lingual Gyrus | 17 |
| L | Caudate |  | L | Anterior Cingulate Gyrus | 24 |
|  |  |  | L | *Cerebellum* |  |
| R | Middle Frontal Gyrus | 9 | R | *Inferior Frontal Gyrus* | 44 |
| R | Superior Frontal Gyrus | 8 | R | Supplementary Motor Area | 6 |
| R | *Inferior Frontal Gyrus* | 45 | R | Precuneus | 7 |
| R | *Inferior Frontal Gyrus* | 44 | R | Fusiform Gyrus | 19 |
| R | Superior Temporal Gyrus | 22 | R | *Cuneus* | 17 |
| R | Middle Occipital Gyrus | 19 | R | *Insula* |  |
| R | *Cuneus* | 18 |  |  |  |

Note: Bilateral results are highlighted. Between-analysis clusters are italicized. The data type is indicated in parentheses.

While small, the total number of studies analyzed here is similar to that found in previous meta-analyses: three of four tasks contrasts in Indefrey (2006) contain between 4-5 studies, Sebastian et al.’s (2011) low and high proficiency groups contained 6 and 8 studies respectively, and Luk et al.’s (2012) language switching metaanalysis included 10 studies. What is more surprising is that, despite the large increase in bilingual neuroimaging studies now available, this increase is only reflected when averaging across a large number of factors, such as in Liu & Cao (2016), who obtained between 17 and 25 studies per contrast by aggregating across task categories. Cargnelutti et al. (2019) boasts up to 37 studies for late L2 learners; however, this is achieved by eliminating all distinguishing factors beyond the presence of L1 or L2 data of any type. When the research question considers very specific or multiple factors, it becomes clear that the relevant data set has hardly grown in fifteen years. Cargnelutti et al. (2019) appears to have just 4 studies from proficient early bilinguals that yield L1>L2 data, and Sulpizio et al.’s (2020) contrasts for grammar and phonology contain just 6 and 5-10 studies, respectively. It is only the very broadly defined lexico-semantics domain and language switching tasks that comprise a data set within the parameters specified by Eickhoff et al. (2016). The minimum number of recommended studies for inclusion in an ALE meta-analysis will be discussed in the data analysis section.

### 2.2 Classification

In the classification of experimental tasks, priority was assigned to the behavioral response each task required from the participant in terms of their perception of and response to stimuli. This necessitated careful reading of each study accepted into the meta-analysis, because often our criteria differed from the manner in which authors described their task. A salient point learned in this process was that the necessary information about task, participant characteristics, and study design is often not labeled clearly or intuitively, which makes the correct and consistent classification of papers a non-trivial aspect of the meta-analysis process. The two authors and three research assistants participated in the classification of tasks. These individuals consulted with one another during the process to resolve any concerns.

Only minimal variation was permitted, as necessary, in order to meet our requirements and still compose coherent task categories with a sufficient number of experiments to perform a meta-analysis. For this reason, studies with word and sentence level prompts were subsumed under the same category, as long as the behavior required from the participant remained consistent. The modality of stimuli presentation was held constant across task categories, with the exception of the syntactic judgment task (see Table 1). Given the aim of our meta-analysis to test for the effect of task design on L1 and L2 processing and to minimize the noise generated by extraneous task demands, and also bearing in mind the limitations of the literature, we consider these exceptions to be well-motivated. Whether possible confounds may have arisen from our categorization scheme will be addressed in the discussion section.

The definition of each task category is provided below. In some cases, additional actions were performed as a comprehension check following the experimental task, but we did not consider it relevant to control for aspects of experimental design that was not measured in the data that authors entered into their analyses. A complete description of each data set is provided in Table 7.

**Table 7.**
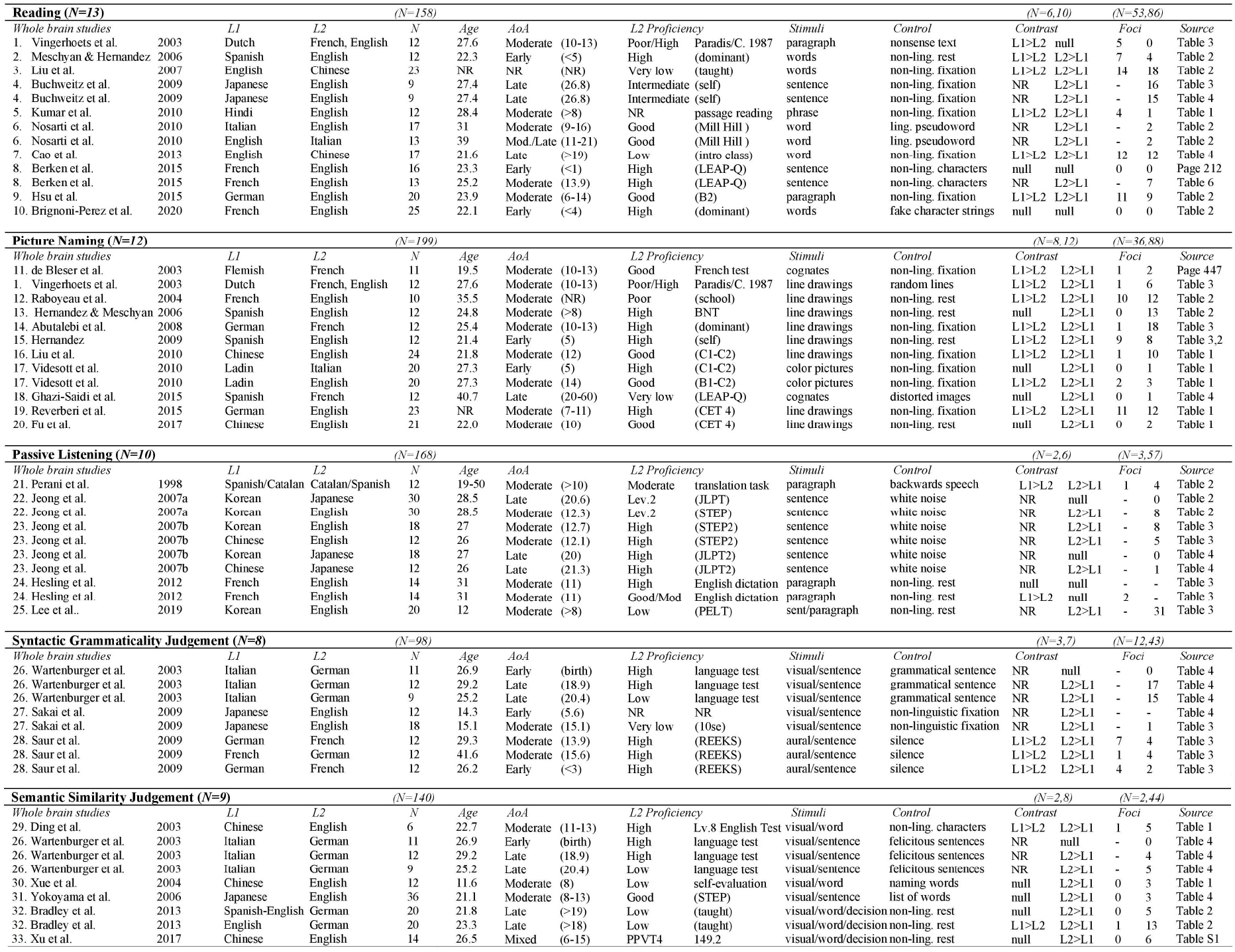
Study Characteristics by Task Type

**Table 8.**
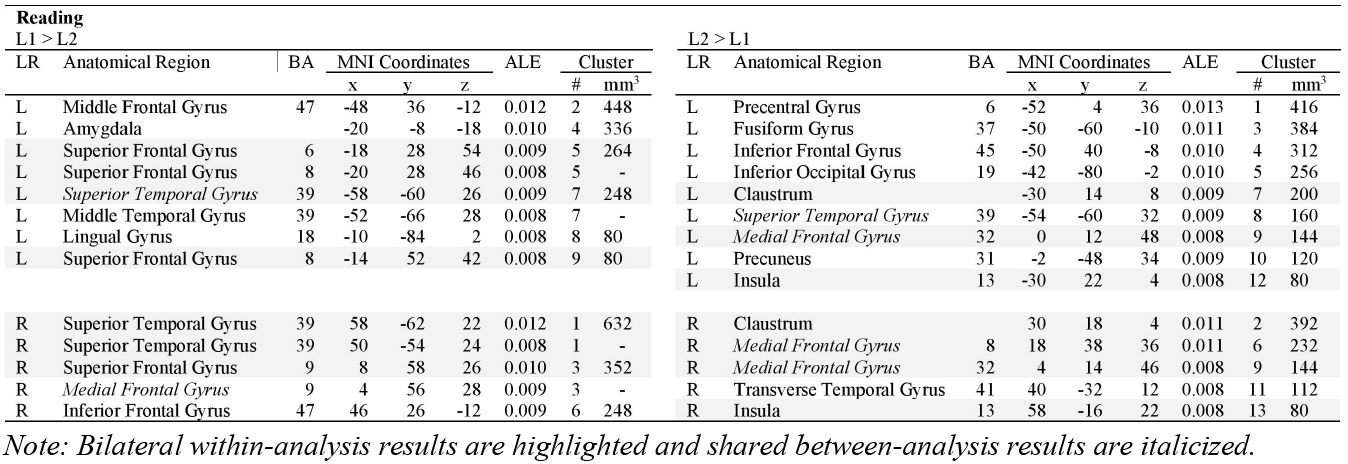
Reading Task: Coordinates

**Table 9.**
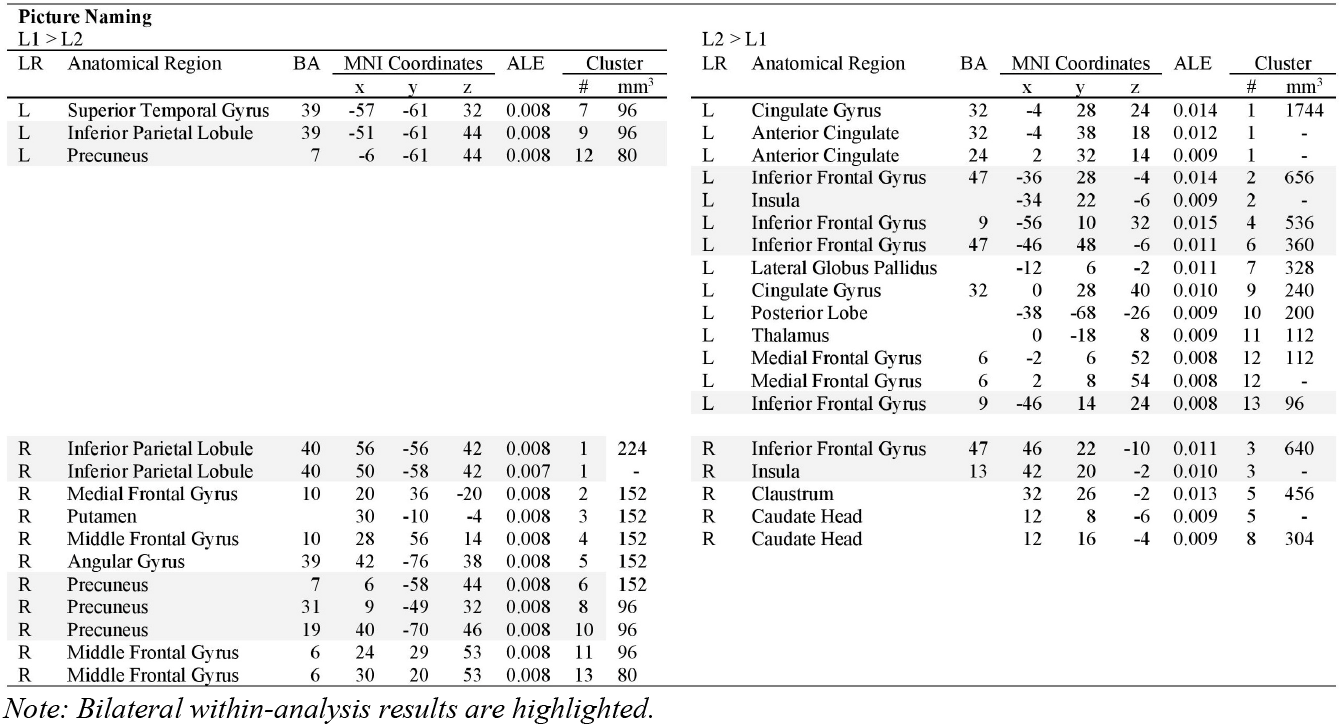
Picture Naming: Coordinates

**Table 10.**
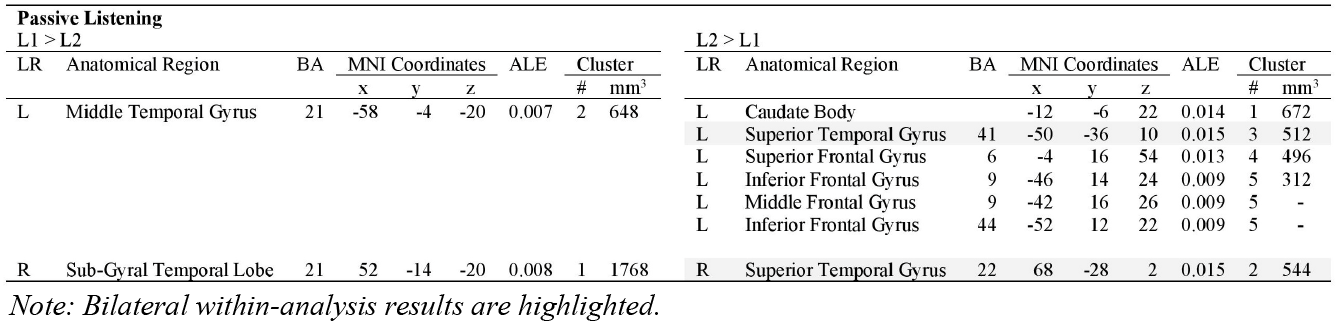
Passive Listening Task: Coordinates

**Table 11.**
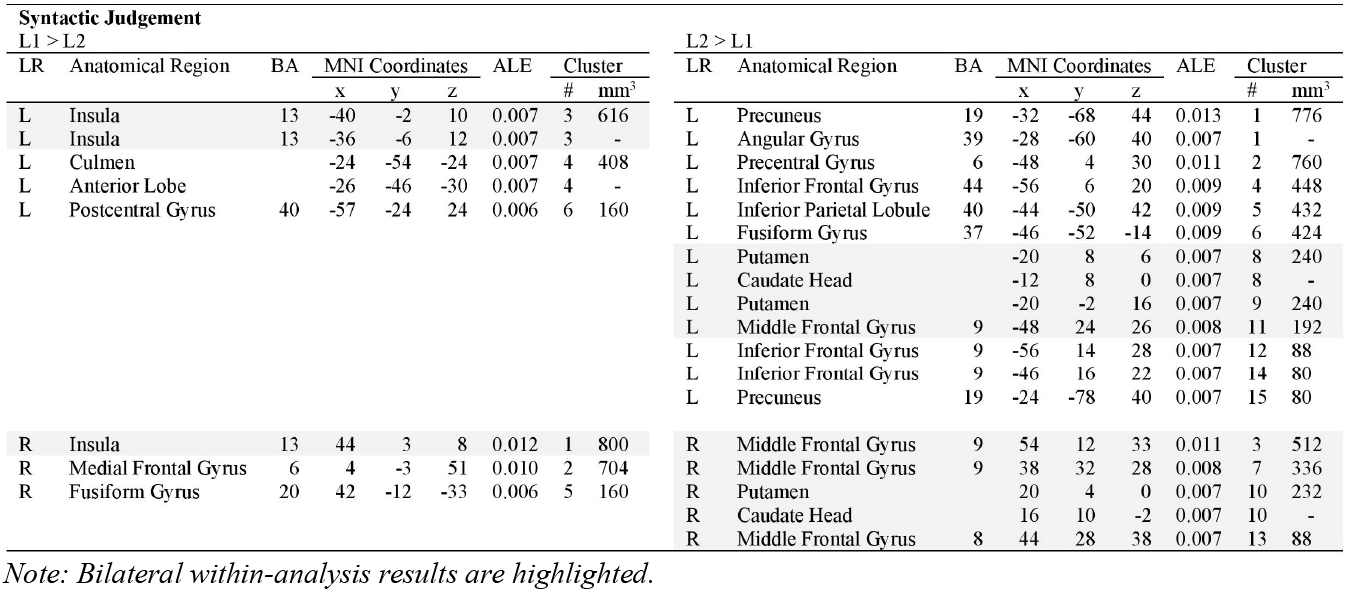
Syntactic Grammaticality Judgment: Coordinates

**Table 12.** Semantic Similarity Judgment: Coordinates

| Semantic Judgement |  |  |  |  |  |  | L2 > L1 |  |  |  |  |  |  |  |  |
| --- | --- | --- | --- | --- | --- | --- | --- | --- | --- | --- | --- | --- | --- | --- | --- |
| L1 > L2 |  |  |  |  |  |  | L2 > L1 |  |  |  |  |  |  |  |  |
| LR | Anatomical Region | BA | MNI Coordinates |  |  | ALE | Cluster | LR | Anatomical Region | BA | MNI Coordinates |  |  | ALE | Cluster |
|  |  |  | x | y | z |  | # mm<sup>3</sup> |  |  |  | x | y | z |  | # mm<sup>3</sup> |
| L | Superior Temporal Gyrus | 39 | -48 | -58 | 22 | 0.008 | 1 744 | L | Angular Gyrus | 39 | -30 | -54 | 38 | 0.011 | 1 1208 |
|  |  |  |  |  |  |  |  | L | Middle Temporal Gyrus | 39 | -30 | -64 | 36 | 0.010 | 1 - |
|  |  |  |  |  |  |  |  | L | Precentral Gyrus | 6 | -42 | 2 | 28 | 0.013 | 2 888 |
|  |  |  |  |  |  |  |  | L | Posterior Cingulate | 29 | -4 | -42 | 16 | 0.012 | 3 800 |
|  |  |  |  |  |  |  |  | L | Middle Frontal Gyrus | 9 | -38 | 22 | 24 | 0.013 | 4 496 |
|  |  |  |  |  |  |  |  | L | Cingulate Gyrus | 23 | 0 | -26 | 28 | 0.012 | 6 432 |
|  |  |  |  |  |  |  |  | L | Precentral Gyrus | 6 | -54 | 2 | 42 | 0.010 | 7 344 |
|  |  |  |  |  |  |  |  | L | Precentral Gyrus | 6 | -56 | 2 | 28 | 0.009 | 8 232 |
|  |  |  |  |  |  |  |  | L | Declive |  | -32 | -80 | -22 | 0.009 | 9 112 |
|  |  |  |  |  |  |  |  | L | Inferior Occipital Gyrus | 18 | -28 | -96 | -6 | 0.009 | 10 112 |
|  |  |  |  |  |  |  |  | L | Region not identified |  | -4 | -88 | -28 | 0.008 | 11 80 |
|  |  |  |  |  |  |  |  | L | Red Nucleus |  | -6 | -25 | -14 | 0.008 | 12 80 |
|  |  |  |  |  |  |  |  | L | Precuneus | 7 | -12 | -66 | 34 | 0.008 | 15 80 |
| R | Precentral Gyrus | 6 | 38 | -10 | 60 | 0.004 | 2 648 | R | Claustrium |  | 32 | 22 | -8 | 0.010 | 5 472 |
| R | Middle Temporal Gyrus | 21 | 64 | -6 | -14 | 0.004 | 3 640 | R | Claustrium |  | 32 | 18 | -4 | 0.009 | 5 - |
|  |  |  |  |  |  |  |  | R | Caudate Body |  | 12 | 8 | 6 | 0.008 | 13 80 |
|  |  |  |  |  |  |  |  | R | Insula | 13 | 36 | 22 | 10 | 0.008 | 14 80 |

#### Reading tasks

These experiments involved covert reading of passages, sentences, or individual words presented visually, with or without probe tasks. Some experiments included reading in multiple orthographies of the same language; if these studies included multiple L1>L2 contrasts with these different orthographies, they were counted as multiple experiments for the purposes of our analysis.

#### Picture naming

These experiments consisted of covert or overt naming of visually-presented pictures, which were either line drawings or color pictures.

#### Passive listening

These experiments involved listening to sentences or paragraphs presented aurally. In order to ensure their attention, participants were asked to respond to comprehension questions after task completion.

#### Semantic similarity judgment

These experiments required a yes or no button-press response to word-level semantic characterization tasks (e.g., “living” or “nonliving”) and sentence-level semantic plausibility judgements (e.g., “the deer shot the hunter” vs. “the hunter shot the deer”). Prompts were presented visually. The sorting of stimuli into one of two categories in lieu of a button-press response was considered to be an association task. There was an insufficient number of association tasks that fit the study criteria for inclusion.

#### Syntactic grammaticality judgment

These tasks required a yes or no button-press response to word-level or sentence-level grammaticality judgements. Visually and aurally presented prompts included violations of agreement (number, gender, case) or word order and verbs with missing arguments (e.g., “he put _ there”).

Additional factors that bear relevance for the analyses are defined below.

#### Baseline conditions

Variation in the baseline control conditions was not considered a meaningful category for exclusion due to our decision to omit data from L1>baseline or L2>baseline contrasts. By utilizing only L1>L2 and L2>L1 contrast data, this factor was eliminated as a potential cause of noise or variation in the results.

#### Proficiency level

The study authors measured participant proficiency level through methods that varied from well-known standardized tests to participant own self-evaluation. The heterogenous nature of these ratings did not allow us to reliably conduct our own classification of proficiency level. Therefore, the level identified by study authors has been accepted with note of the different means of measuring proficiency (see Table 7).

#### AoA

The age at which an AoA effect may occur has been fiercely debated in the literature, although the critical period for language acquisition is often set at the age of 5-6 years. We consider recent work by Hartshorne, Tenenbaum, and Pinker (2018), who propose sensitive periods for ultimate language attainment with thresholds at the age of 9 and 16 years. We will compare our results with alternative AoA categories in the discussion section. Studies that did not report AoA were included into the main task analysis and excluded from the AoA analysis.

### 2.3 Data Analysis

The activation likelihood estimation (ALE) method for neuroimaging (Eickhoff, Bzdok, Laird, Roski, Caspers, Zilles, & Fox, 2011; Eickhoff, Bzdok, Laird, Kurth, & Fox, 2012; Eickhoff, et al. 2009; Turkeltaub, Eickhoff, Laird, Fox, Wiener, & Fox, 2012) differs from behavioral meta-analyses in that the location and spatial uncertainty of effects are evaluated, rather than reported effect sizes. The contributing experiments provide the stereotactic coordinates of foci identified in their individual fMRI analyses. These coordinates are then modeled as Gaussian probability distributions that are weighted for the number of subjects in each experiment, and the likelihood of convergence across foci is calculated from the distributions as the above-chance probability of clustering between experiments in a random-effects analysis. Standard *p* values are used to measure whether the likelihood of convergence is greater than predicted under a null distribution of random spatial association, and the ALE score represents the relative convergence of foci. Because *p* values are set to a predetermined level prior to the analysis to limit the output, ALE scores representing the greater spatial consistency of a cluster are typically reported with the size of the estimated cluster. However, the number of experiments that target the “true” location of an effect may vary considerably, which becomes a risk in small datasets and may affect the reliability of the results.

While careful to note that the optimal approach depends on the research question and the characteristics of the data set, the authors recommend performing cluster-level family-wise error (FWE) correction with the inclusion of no less than 17-20 studies, which limits the contribution of the most dominant experiments to less than 50% and the contribution of the top two experiments to less than 80% (p 74). Voxel-level family-wise error correction is also considered rigorous and may require only eight studies; however, this analysis results in substantially smaller cluster sizes and is often too stringent a requirement for most meta-analyses. Particularly harsh criticism was reserved for voxel-level false discovery rate (FDR) correction (Laird, Fox, Price, Glahn, Uecker, Lancaster, Turkeltaub, Kochunov, & Fox, 2005; Wager, Lindquist, & Kaplan, 2007), which was deemed generally unacceptable for meta-analyses. This method may lead to false negative and false positive findings simultaneously (Chumbley & Friston, 2009; Eickhoff et al., 2009; Genovese, Lazar, & Nichols, 2002) because the number of incidental clusters generated grows together with the number of experiments that identify the true effect. The authors also warn against an uncorrected threshold, which is typically set at *p* < .001 with an arbitrary additional extent threshold. While uncorrected thresholding was the most sensitive to true effects and displayed the greatest power, uncorrected thresholding was also more prone to reveal spurious findings with results that were likely to be driven by the most dominant experiment. Generally, 28 experiments are recommended for uncorrected thresholding at *p* < .001 to ensure that the contribution of the experiment with the greatest number of participants does not exceed 50% (Eickhoff et al., 2016).

Out of necessity, the existing ALE meta-analyses have not been able to fully comply with these recommended best practices. In addition to the aforementioned limitations on the number of available experiments, only Cargnelutti et al. (2019) employs FWE correction for their main effect analyses. Sebastian et al. (2011) and Liu & Cao (2016) utilize FDR correction (*p* < .05) with a minimum cluster size of 150mm^3^, as does Luk et al. (2012), with a lower threshold *(p* < .01) and a minimum cluster size of 100mm^3^. Contrast analyses in Liu & Cao (2016) and Cargnelutti et al. (2019) were carried out with uncorrected thresholding: the former at *p* < 0.05 and a minimum cluster volume of 150mm^3^ and the latter at *p* < .001 with a minimum cluster size of 80mm^3^. Liu & Cao (2016) took additional measures to only include foci reported in no less than two studies when the total number was < 15, or no less than three studies, when the total number was > 15 (cf. Turkeltaub & Branch Coslett, 2010; Wagner, Sebastian, Lieb, Tuscher, & Tadic, 2014). Yet this measure is unlikely to counteract the additional concerns raised about FDR correction. Sulpizio et al. (2020) conducted all their analyses at uncorrected thresholding with *p* < .001 and a minimum cluster size of 150mm^3^.

Unfortunately, the data set for our task-based meta-analysis also remains too small to fully comply with all of Eickhoff et al.’s (2016) recommendations for best practice, but we have endeavored to adopt appropriate guidelines that are comparable to those employed in previous bilingual ALE meta-analyses. We have chosen to use uncorrected thresholding at *p* < .001 with a minimum cluster size of 80mm^3^ for several reasons. This method was utilized for the contrast analyses in the three most recent meta-analyses, allowing a clear basis of comparison with the established literature. Moreover, the spurious clusters that arise in meta-analyses are thought to be caused by noise generated due to heterogenous task demands (pp. 78, 81) and the variation that is unavoidable in a larger data set of studies: Eickhoff et al. (2016) caution that strong effects which would be present in the majority of experiments found within a typical ALE meta-analysis data set are assumed to be rare (p. 78). Therefore, our strategy to utilize only the contrasts between L1 and L2 and to include less studies, but those with greater similarity in task design should minimize noise and maximize the consistency of real effects in our data set, allowing us to capitalize on the greater sensitivity to true effects obtained through uncorrected thresholding and yield findings that may be even more reliable than those obtained in larger meta-analyses with heterogenous tasks.

Additionally, our meta-analysis provides a novel contribution to the existing literature by investigating how many foci each study contributes to the clusters generated by each contrast. This is a measure of how consistent each study may be in producing the overall findings and how reliably the findings may reflect the underlying population of experiments. In order to visualize this information, foci from each contributing study were overlaid on ALE maps and color-coded. In the results and discussion sections, we will consider which studies provided the most relevant findings and whether the most heavily weighted studies from each data set played a disproportionate role, as cautioned by Eickhoff et al. (2016), or whether disparities in the contribution of each experiment may be related to experimental design features.

The following steps were taken to perform the ALE meta-analyses and create the figures included in this paper. The relevant foci reported in the contributing studies were extracted. Coordinates reported in Talairach space were converted into Montreal Neurological Institute (MNI) standard space by the icbm2tal transform (Laird, Robinson, McMillan, Tordesillas-Gutierrez, Moran, Gonzales, Ray, Franklin, Glahn, Fox, & Lancaster, 2010; Lancaster, Tordesillas-Gutierrez, Martinez, Salinas, Evans, Zilles, Mazziotta, & Fox, 2007). ALE analyses were implemented in GingerALE 3.0.2 (http://www.brainmap.org/ale). Contrasts between L1>L2 and L2>L1 were performed within each task subset using a threshold at *p* < .001 (uncorrected) with 1,000 permutations and a minimum cluster volume of 80mm^3^ (the same cluster extend threshold selected for contrast analyses by Cargnelutti et al., 2019; Liu & Cao, 2016). The thresholded ALE maps were overlaid onto the MNI152 coordinate brain template using the MRIcroGL software (https://www.nitrc.org/projects/mricrogl). For task-specific maps, foci were located according to their stereotactic coordinates and manually marked onto the map in FSLeyes (FMRIB’s Software Library, www.fmrib.ox.ac.uk/fsl). Image editing software was then utilized to render the foci as dots on the final maps, identifying the foci from each experiment according to a unique color.

## 3. Results

When combining across tasks, our meta-analysis results closely parallel those found in a recent highly powered meta-analysis (Cargnelutti et al. 2019). The use of a smaller data set seems not to have generated spurious clusters, and the map of combined tasks illustrates how the L1 and L2 networks may have differentially contributed to the earlier results. When individual tasks are disambiguated, we observe activation that still remains within the general outline of the language network, although somewhat more distributed. Clusters in areas like the cerebellum, which have been reported in numerous individual studies, are revealed to be more common to particular experimental paradigms. Importantly, the L1 and L2 networks display non-overlapping fMRI activation by task type. These findings indicate we may legitimately propose that different linguistic competencies recruit certain areas of the language network to a greater or lesser degree in the L1 and L2 networks. In particular, we see evidence that the L2 network is more focal, whereas the L1 network appears to activate regions peripheral to core language areas.

### 3.1 Reading

**Overall**, reading experiments revealed a distributed network in language areas to which many of the studies that make up the data set contribute. The L2 network revealed greater engagement of morphological processes, with the possible exception of syntactic judgment, and relatively little subcortical fMRI activation.

The **L1>L2 contrast** found substantial bilateral fMRI activation within similar regions in each hemisphere. Of these two clusters, there is more activation of the right temporal lobe. <u>Bilateral activation</u> occurred in the *superior frontal gyrus* (left BA 6/8, right BA 9) and the *superior temporal gyrus* (BA 39). In the <u>left hemisphere</u>, three additional clusters were found in frontal regions in the *middle frontal gyrus* (BA 47), in temporal regions in the *middle temporal gyrus* (BA 39), and the *lingual gyrus* (BA 18) of the occipital lobe. Considerable subcortical activation was found in the *amygdala*. In the <u>right hemisphere</u>, additional activation was found in frontal regions: extending into the *medial frontal gyrus* (BA 9) and in one new cluster in the *inferior frontal gyrus* (BA 47).

The **L2>L1 contrast** also revealed substantial bilateral fMRI activation, but in three clusters located in different regions. <u>Bilateral activation</u> occurred outside core language areas in the *claustrum* and *insula* (BA 13), as well as in the *medial frontal gyrus*, particularly in the right hemisphere (left/right BA 32, right 8). Additional significant activation was concentrated in the <u>left hemisphere</u>, with larger clusters found in posterior regions. This included five clusters in the *precentral gyrus* (BA 6), *fusiform gyrus* (BA 37), *inferior occipital gyrus* (BA 19), *superior temporal gyrus* (BA 39), and *precuneus* (BA 31). Frontal regions produced one cluster in the *inferior frontal gyrus* (BA 45). In the <u>right hemisphere</u>, just one additional cluster was found in the *transverse temporal gyrus* (BA 41).

The reading task forms one of the larger data sets, yet there is low consistency in the location of foci reported from each study. This may be unsurprising, considering the numerous subprocesses involved in reading and the absence of a shared response strategy in experiments, which might have served to orient participants and engage similar preparatory subprocesses. The large number of foci positioned in proximity to a cluster may indicate where a larger data set could elicit additional or more extensive clusters. Nonetheless, the evidence does not suggest that the results are driven by the most heavily weighted experiments—those with the largest number of participants.

In the **L2 network**, five out of thirteen clusters are generated by foci from at least two experiments, with contributions by all studies in the data set. The most heavily weighted experiment (Liu et al., 2007) shares four of six clusters with foci from other studies, and the second ranked experiment (Hsu et al., 2015) does not fall within any cluster, although it contributes foci that lie adjacent to clusters. Buchweitz et al. (2009) and Cao et al. (2013) contribute five and four clusters, respectively. In the **L1 network**, the results are more strongly influenced by the most heavily weighted study. Liu et al. (2007) produces seven clusters, six of which are shared with other studies. The second ranked study (Hsu et al., 2015) shares foci with other studies in five out of six clusters.

### 3.2 Picture Naming

**Overall**, picture naming experiments revealed an L2 network highly concentrated in left frontal and subcortical regions and a widely distributed and right-lateralized L1 network. This task provides particularly strong evidence for a disassociation between the networks, particularly given that it possesses the largest L1 and L2 data sets.

The **L1>L2 contrast** produced ten of thirteen clusters in the right hemisphere. <u>Bilateral activation</u> was found within a limited posterior region, in the *inferior parietal lobule* (left BA 39, right BA 40) and *precuneus* (left BA 7, right BA 7/19/31). In the <u>left hemisphere</u>, scattered foci occur in frontal areas and the cerebellum, but only one additional cluster was generated in the *superior temporal gyrus* (BA 39). In the <u>right hemisphere</u>, two additional clusters were found in a subcortical region, the *putamen*, and adjacent to the bilateral cluster in the *angular gyrus* (BA 39). Four additional clusters were found in frontal regions in the *medial* and *middle frontal gyrus* (BA 6/10).

The **L2>L1 contrast** revealed a left-lateralized network. <u>Bilateral activation</u> appeared in the *inferior frontal gyrus* (BA 47) and subcortically in the *insula* (right BA 13). In the <u>left hemisphere</u>, six additional clusters occurred, primarily in subcortical areas: the *cingulate gyrus* (BA 9/32), *anterior cingulate* (BA 32), *lateral globus pallidus*, and *thalamus* (BA 11). Two clusters were found in the cerebellar posterior lobe and *medial frontal gyrus* (BA 6). In the <u>right hemisphere</u>, clusters arose in the *claustrum*, an inner cortical area, and the subcortical *caudate head*.

Despite the larger data set in this task, the **L1 network** still appears to be influenced by the two most heavily weighted studies to a greater degree than the **L2 network** does. Three of the eight experiments from the **L1 network** have nearly twice as many subjects. Liu et al. (2010) contributes one foci and cluster, whereas Reverberi et al. (2015) generates all but two other clusters. Videsott et al. (2010) contributes in the *inferior frontal gyrus*. The foci in the L1 data set are widely scattered with little consistency in their location. In the **L2 network**, roughly half of the experiments have at least twenty subjects and half of clusters are generated by two or more studies. Liu et al. (2010) contributes to four of the thirteen clusters, or less than a third of the total. Reverberi et al. (2015) also contributes to half of the clusters. All the studies in this contrast data set contribute to at least one cluster.

### 3.3 Passive Listening

**Overall**, passive listening experiments elicited a smaller number of clusters than any other task and no fMRI activation was shared between the L1 and L2 networks. Passive tasks have typically produced a less substantial hemodynamic response than active ones, which could hinder identification of the full range of each network.

The **L1>L2 contrast** generated two clusters within the temporal lobe. <u>Bilateral activation</u> occurred in the left *middle temporal gyrus* (BA 21) and more extensively in the right *sub-gyral temporal lobe* (BA 21).

The **L2>L1 contrast** also illustrated bilateral activation within the temporal lobe, but somewhat higher than in the first contrast, in the *superior temporal gyrus* (left BA 41, right BA 22). The activation in the <u>right hemisphere</u> was more extensive for this contrast as well. In the <u>left hemisphere</u>, three additional clusters were found in frontal regions in the *superior frontal gyrus* (BA 6), *inferior frontal gyrus* (BA 9/44), and *middle frontal gyrus* (BA 9). The largest cluster in this network was found subcortically in the *caudate body*.

In the **L1 network**, the data set comprised experiments conducted by just two sets of authors. The experiments contributed three foci, all of which generated clusters. However, in the right hemisphere, the two studies appear consistent, as they both contribute to the same cluster in the *temporal lobe*. A unique aspect of this task data set was that one set of authors (Jeong et al., 2007) contributed four experiments to the total data set of six experiments that describe the **L2 network**. The fact that other authors had difficulty eliciting the same result may indicate that differences in bilingual language processing are less common in passive listening tasks than in other linguistic competencies, or that Jeong et al. (2007) has some unidentified, unique element to their experimental paradigm. However, according to the general assumptions of our task-based meta-analysis, we should in fact expect substantial overlap in the location of foci from these experiments, because the data points were generated by identical experimental paradigms. Three of the experiments contributed between five and eight foci each, and three to four foci per experiment occur in close proximity within the same clusters. The fourth experiment provided just one focus, which did not generate a cluster, but occurred nearby to two foci in the *superior temporal lobe* from Lee et al. (2019). Somewhat surprisingly, Lee et al. (2019) contributed no less than thirty-one foci, substantially more than any other study. These foci were widely distributed across the exterior and subcortical regions of both hemispheres. Three of the foci appeared near clusters in the frontal and temporal lobes and two directly contributed to the cluster in the left *caudate body*. The final remaining study, Perani et al. (1998), contributed four foci, all in the right hemisphere and all located at some distance from the foci produced by other studies. Two of these were in the cerebellum, and two in posterior regions.

### 3.4 Syntactic Grammaticality Judgment

**Overall**, syntactic judgment experiments revealed a largely left-lateralized L2 network with substantial fMRI activation in motor and posterior regions and a more bilateral L1 network. The two networks shared no common clusters, although frontal and subcortical activation occurs in neighboring regions.

The **L1>L2 contrast** produced bilateral activation that was somewhat more extensive in the right hemisphere. The largest clusters showed <u>bilateral activation</u> in the bilateral *insula*. In the <u>left hemisphere</u>, two additional clusters were generated in the cerebellar *culmen* and *anterior lobe* and in the *postcentral gyrus* (BA 40). In the <u>right hemisphere</u>, two additional cluster arose in the *medial frontal gyrus* (BA 6) and *fusiform gyrus* (BA 20).

The **L2>L1 contrast** generated three bilateral clusters and eight in the left hemisphere. No clusters were unique to the right hemisphere. <u>Bilateral activation</u> included clusters in the subcortical *putamen* and *caudate head* and the *middle frontal gyrus* (left BA 8, right BA 8/9). In the <u>left hemisphere</u>, an additional eight clusters were found, the largest being in posterior regions in the *precuneus* (BA 19) and *angular gyrus* (BA 39). Other posterior clusters were located in the *inferior parietal lobule* (BA 40) and *fusiform gyrus* (BA 37). The *precentral gyrus* (BA 6) of the motor cortex revealed one cluster and the *inferior frontal gyrus* (BA 9/44) produced three clusters.

This task produced results that were possibly the most consistent of all five tasks. All of the foci from the data set appeared within proximity of clusters. To some extent, this maybe unsurprising, given that two sets of authors contributed multiple experiments (Saur et al., 2009; Wartenburger et al., 2003). Only Saur et al. (2009) found results for the **L1 network**, although all of their experiments contributed to L1 clusters. Only clusters in the right *fusiform gyrus* and left *postcentral gyrus* were elicited by a single experiment. In the **L2 network**, the majority of clusters were generated by at least two foci from different studies. The most heavily weighted experiment (Sakai et al., 2009) contributed just one focus and cluster, however this cluster is close in proximity to other foci in the *inferior frontal gyrus* from two separate studies. Four of the five remaining experiments have equal weight, and clusters in frontal, posterior, and subcortical areas are supported by studies from both sets of authors (Saur et al., 2009; Wartenburger et al., 2009) with eleven foci each in frontal regions, and one focus each in the *fusiform gyrus*. Wartenburger et al. (2009) show particularly consistent results. Clusters in the *precuneus, angular gyrus*, and *inferior parietal lobule* were primarily generated by Wartenburger et al. (2009), but Saur et al. (2009) contributed one focus in a cluster in the *precuneus*. Right hemisphere clusters were exclusively generated by Wartenburger et al. (2009).

### 3.5 Semantic Similarity Judgment

**Overall**, the semantic judgment task revealed a left-lateralized L2 network with substantial fMRI activation in motor and posterior regions which closely resembles the network found for the syntactic judgment task. The L1 network, however, is concentrated in the right hemisphere. There is no bilateral activation or shared activation between hemispheres. Some frontal and subcortical clusters are adjacent to clusters in the opposite hemisphere.

The **L1>L2** contrast produced three clusters in the *temporal lobe* and *pre-central gyrus*, with no <u>bilateral activation</u>. In the <u>left hemisphere</u>, activation was found in the *superior temporal gyrus* (BA 39). In the <u>right hemisphere</u>, two clusters were identified in the *precentral gyrus* (BA 6) and the *middle temporal gyrus* (BA 21).

The **L2>L1 contrast** generated twelve clusters with widespread activation across frontal, posterior, and subcortical regions. <u>Bilateral activation</u> was absent. In the <u>left hemisphere</u>, twelve clusters where produced, more than in any other task. The largest of these occurred in posterior regions, in the *angular gyrus* (BA 39) and *middle temporal gyrus* (BA 39), with two smaller clusters in the *precuneus* (BA 7) and *inferior occipital gyrus* (BA 18). An additional three clusters were found in the *precentral gyrus* of the motor cortex (BA 6). Subcortical regions were identified in the *posterior cingulate* (BA 29), *cingulate gyrus* (BA 23), cerebellar *declive*, and the *red nucleus* in the midbrain. One cluster was found in a frontal region, the *middle frontal gyrus* (BA 9). In the <u>right hemisphere</u>, activation was found in inner cortical and subcortical regions: the *claustrum*, *insula* (BA 13), and *caudate body*.

This task encompassed a larger number of authors and still results nearly as consistent as the syntactic grammaticality judgment task. Foci in the left hemisphere that did not generate clusters were located in areas similar to those found in the other decision task. Clusters generally contained at least two foci from different experiments, particularly in the left hemisphere. In the **L1 network**, three foci all generated their own cluster in widely dispersed regions. The most heavily-weighted study (Bradley et al., 2013) produced the left hemisphere cluster, and a study with just six subjects (Ding et al., 2003) generated the two right hemisphere clusters. In the **L2 network**, the largest cluster involved four of the eight total experiments, and four experiments produced two clusters in the motor cortex. The most heavily weighted study (Yokoyama et al., 2006) contributed three isolated foci in the *occipital lobe* and *cerebellum*. Two experiments from Bradley et al. (2006) were ranked second and contributed thirteen and five foci, respectively. Both produced clusters in the *middle frontal gyrus* and subcortical *cingulate gyrus*. An additional two clusters in the *anterior cingulate* and *angular gyrus* and *middle temporal gyrus* were generated in conjunction with other authors. Foci from Wartenburger et al. (2003) and Bradley et al. (2013) appeared in subcortical areas. Bradley et al. (2013) contributed two foci in a morphological processing areas.

## 4. Discussion

Meta-analyses are intended to provide a reliable average of the relevant fMRI activation obtained for a particular research question by virtue of large data sets that are composed of studies conducted in multiple labs. However, the choice of what data to include and individual differences inherent in the participant population may still substantially affect the individual neuroimaging studies (e.g., Blank, Balewski, Mahowald, & Fedorenko, 2016) that are used as inputs for a meta-analysis, and therefore the results of the meta-analysis itself. For this reason, we will consider what results appear to be consistent across meta-analyses that divide up language tasks in different ways, and we will note not only when there is a direct replication of activation found in another study, but when activation shares either the same Brodmann area or anatomical region and lies adjacent to the actual cluster identified. Finally, we will consider how variation in the tasks included in other meta-analyses may have influenced their findings, as well as how the characteristics that define the most heavily weighted studies in each linguistic task analysis may have influenced our own findings.

### 4.1 Reading

Previous meta-analyses that differentiate linguistic competencies lack a category that would exclusively represent reading tasks. In Indefrey’s (2006) sentence-level listening and reading category, only three out of fourteen experiments employ reading tasks, and of these, none contributed to the group level findings. Therefore, it may be that the three clusters identified in left frontal regions of the L2 network by Indefrey (2006) are representative of sentence-level rather than reading-specific processes. The reading task analysis revealed a network with greater engagement of posterior regions, and clusters in frontal regions were more anterior (BA 45), interior (BA 32), and bilateral (BA 32). The greater similarity between these results and those generated by Sulpizio et al.’s (2020) lexico-semantic domain may stem from the more extensive activation generated in the later study. Although only a small fraction of the data set included reading tasks (three of eighteen studies in the L1 network and four of twenty-two studies in the L2 network), reading involves a large number of subprocesses that could possibly have been elicited by other experimental designs in this heterogenous collection of studies. In the L1 network, the lexico-semantic data set revealed greater posterior and subcortical activation than the reading analysis, which produced greater bilateral *temporal lobe* activation. In the L2 network, the reading task generated the more expansive network, incorporating posterior regions and bilateral activation in fontal and temporal regions. Both analyses generated clusters in the bilateral *insulae* (BA 13) and substantial subcortical activation, although this was centered in the bilateral *pallidum* for lexico-semantic tasks and the right *caudate* for reading tasks.

Indefrey’s (2006) and Sulpizio et al.’s (2020) results overlap partially with a recent meta-analysis devoted entirely to bilingual reading tasks (Li, Zhang, & Ding, 2021). The forty studies subsumed under this category by the authors are still quite heterogenous, including tasks such as homophone matching, sound judgment, and one-back matching tasks, among others. In the L1 network, Sulpizio et al. (2020) identified several clusters in left frontal regions that are adjacent to those found in Li et al. (2021), yet they match poorly with the activation generated by the reading task, which was located somewhat lower in left frontal (BA 6, BA 47) and posterior regions (BA 37, BA 18). No common activation was found in the right hemisphere. In the L2 network, Indefrey’s (2006) sentence-level analysis replicates two clusters in the left *supplementary motor area* (BA 6) and *middle frontal gyrus* (BA 9), and Sulpizio et al.’s (2020) lexico-semantic domain replicates a cluster in the left *insula* (BA 13). However, the frontal activation found in Sulpizio et al. (2020) is largely absent in the later study, and most of the bilateral posterior activation in Li et al. (2021) appears unique. The reading analysis replicated a left hemisphere cluster in the *insula* (BA 13) and *fusiform gyrus* (BA 37), and provided adjacent clusters in frontal (BA 6) and posterior regions (BA 19). No activation was shared in the right hemisphere. The more stringent L2>L1 contrast in Li et al. (2021) again replicated the left *insula* (BA 13) in the L2 network and generated an adjacent cluster (BA 39).

Overall, the reading task elicited similar L1 and L2 networks characterized by temporal and inner or inferior frontal activation. Greater subcortical activation was found in the L2 network. Li et al. (2021) proved most similar in its findings, generating similar L1 and L2 networks with left hemisphere clusters high in frontal and posterior regions. Activation unique to each network fell within the areas noted in our analysis. The results of Indefrey (2006) are compatible but incomplete, whereas Sulpizio et al.’s (2020) findings appear largely inconsistent with the reading task results. Moreover, little engagement of the language control network was evident, although clusters adjacent to Luk et al.’s (2012) findings were identified in both networks in frontal and temporal regions.

When considering proficiency level, the two most heavily weighted studies in the L1 network featured low proficiency individuals, who predominated throughout the data set. A wider range of proficiency levels were represented in the data set for the L2 network, although more foci were contributed by low proficiency bilinguals. In Sebastian et al.’s (2011) proficiency level analysis, the L1 network is largely compatible with clusters in bilateral temporal and left posterior regions adjacent to those from the reading task (BA 39, BA 18). In the L2 network, left hemisphere activation is more localized in frontal and subcortical regions than in the reading task, although adjacent clusters appear in the reading task (BA 45, BA 32, BA 6). The task replicates clusters in the left *precuneus* within a different Brodmann area (BA 31) and subcortically in the right *claustrum*. No results from this high proficiency group were replicated in either network of the reading analysis, although one adjacent cluster was found in posterior regions (BA 18). Cargnelutti et al.’s (2019) meta-analysis only considered highly proficient bilinguals at an early and late AoA. Very minimal results were obtained for the L1 network of both groups and only the left *middle temporal gyrus* (39) for early AoA replicated in the reading task. In the L2 network, highly proficient bilinguals with late AoA generated a wide range of clusters that replicated in the bilateral *insulae* (BA 13) and fell within regions adjacent to our reading task in the left hemisphere (BA 45, BA 32, BA 39). Key differences included more widespread activation in the right hemisphere for the reading task in frontal and temporal regions and the earlier meta-analysis found only subcortical right hemisphere activation.

When considering AoA, the two most heavily weighted studies in the L1 network featured participants with moderate to late AoA. In the L2 network, the majority of the data set including the most heavily weighted studies comprised bilinguals with moderate AoA. However, the AoA analyses performed by Liu & Cao (2016) overlapped considerably between AoA categories, with findings localized in frontal and subcortical regions and without right hemisphere activation. In the L1 network, the reading task replicated a cluster in the left *precentral gyrus* (BA 6) and generated an adjacent cluster the left *temporal lobe* (BA 39) for both early and late AoA. In the L2 network, these clusters in the left *precentral gyrus* (early BA 6, late BA 44) and *insula* (early BA 13, late BA 44) are again replicated or found adjacent, without temporal activation. The primary difference between early and late bilinguals is in the expanded scope of activation found for late bilinguals, and the more specific reading task produced greater bilateral and right hemisphere results in both networks. Cargnelutti et al.’s (2019) AoA analyses generated the same approximate scope of activation as the earlier study, with more right hemisphere involvement. For both networks, early bilinguals generated clusters that fell within the activation produced by late bilinguals. In the L1 network, the only cluster to replicate was the *middle temporal gyrus* (BA 39) for both groups. Clusters adjacent to those in the reading task were found in frontal (BA 6) regions for both AoA and posterior regions (BA 18) for late AoA. In the L2 network, both early and late AoA replicated clusters in the left *inferior frontal gyrus* (BA 45), *medial frontal gyrus* (BA 32), *superior temporal gyrus* (BA 39) and generated an adjacent cluster in the *precuneus* (BA 31). Late bilinguals replicated activation in the bilateral *insulae* (BA 13).

Therefore, in considering what effect participant characteristics may have on the analysis, it appears that during reading tasks, <u>low proficiency bilinguals</u> engage bilateral temporal areas and a wider range of posterior regions in their **L1 network**, particularly when they are late bilinguals. The **L2 network** of low proficiency bilinguals may be relatively more widespread, with greater bilateral activation in frontal, posterior, and subcortical regions. <u>High proficiency bilinguals</u> tend to engage motor and posterior regions in the **L1 network**, whereas in their **L2 network**, more frontal areas are involved. Late bilinguals again generate a wider scope of activation, particularly in posterior regions. Right hemisphere activation appears to be less related to AoA and more to proficiency level.

### 4.2 Picture Naming

The picture naming task is analyzed as an independent linguistic competency in Indefrey (2006). However, this category generates just one cluster in the left *posterior* and *inferior frontal gyrus* (BA 44/47) of the L2 network. Our picture naming analysis replicates this finding in two clusters (BA 47), which is unsurprising as the area represents an important region of the classical language network. In Sulpizio et al.’s (2020) lexico-semantic domain, picture naming tasks make up nearly half of the data set: eight out of eighteen (L1) and twenty-two (L2) experiments, including half of the most highly weighted studies. In the L1 network, clusters in the left *precuneus* (BA 7) and right *angular gyrus* (BA 39) replicate in the task analysis and others are found in adjacent temporal (left BA 39) and posterior regions (right BA 40). Considerably more right hemisphere activation is found in the task analysis, whereas the lexico-semantic domain generates a response that is more balanced between frontal and posterior regions, particularly in the left hemisphere. In the L2 network, only the left *inferior frontal gyrus* (BA 47) and bilateral *insula* replicate, although there substantial frontal activation (BA 9, BA 6) in adjacent clusters in the left hemisphere are noted. In the right hemisphere, both analyses show primarily inner or subcortical activation, whereas the picture naming task generates a much wider network in the left hemisphere.

Overall, it can be concluded that despite the task forming nearly half of the data set, Sulpizio et al.’s (2020) lexico-semantic domain poorly represents the picture naming and Indefrey’s (2006) analysis is insufficient in scope. Picture naming appears to be characterized by greater engagement of posterior and subcortical regions in the L1 network, particularly in the right hemisphere. In the L2 network, picture naming strongly engages the left hemisphere, primarily in subcortical regions and to a somewhat lesser degree in frontal regions. Picture naming exhibits greater engagement of the language control network than reading, but primarily in the L2 network. This is seen in the *caudate head* and areas adjacent to Luk et al.’s (2012) findings in frontal regions (BA 6, BA 9/46, BA 47). In the L1 network, the *middle frontal gyrus* (BA 6) rather than the *precentral gyrus* is engaged.

When considering proficiency level, the data set for picture naming almost entirely comprised participants with a high proficiency level. One study with low proficiency participants contributed very few foci, and the other contributed numerous foci that resulted in very few clusters. Nonetheless, Sebastian et al.’s (2011) analysis of high proficiency participants poorly approximates the picture naming results. In the L2 network, no clusters are replicated, although clusters adjacent to those in the task analysis are present in left frontal regions (BA 46, BA 9). In the L1 network, the high proficiency analysis predominately found left-lateralized activation in posterior regions, whereas the picture naming task elicited a wide range of right hemisphere activation. In the left hemisphere, picture naming replicated a cluster in the *precuneus* (BA 7). Greater overlap between the two analyses was seen in the L2 network for low proficiency individuals. Clusters in the left hemisphere replicated in the *inferior frontal gyrus* (BA 47), *cingulate* and *paracingulate gyrus* (BA 32, BA 24), and adjacent clusters were found in frontal regions of both hemispheres (left BA 9, right BA 47). In the L1 network, a cluster adjacent to those in the task analysis was found in the left *temporal lobe* (BA 39). In contrast, Cargnelutti et al.’s (2019) high proficiency analysis shared no common activation in the L1 network with the task analysis, whereas clusters in the L2 network replicate in the *inferior frontal gyrus* (BA 47) for early and late AoA. The late AoA findings are extensive, and additional replication is found in the left *medial frontal gyrus* (BA 6), right *inferior frontal gyrus* (BA 47), right *caudate*, and the bilateral *insulae* (BA 13).

When considering AoA, the picture naming data set is composed predominately of participants with moderate AoA, including the two most heavily weighted studies. Liu & Cao’s (2016) early AoA analysis is consistent with the picture naming analysis in the L2 network, replicating clusters in the left *inferior frontal gyrus* (BA 9) and *insula*, and producing adjacent clusters in frontal regions (BA 6, BA 10). For late AoA, the earlier meta-analysis found clusters in left frontal regions adjacent to those of the task analysis (BA 9, BA 8, BA 46, BA 44). The only cluster to replicate is the left *insula*. Liu & Cao (2016) find no right hemisphere activation for any of their AoA contrasts. In the L1 network for early AoA, the earlier study produced frontal clusters that are not present in the task analysis, as well as posterior clusters in adjacent posterior regions (BA 7, BA 39). The late AoA group differs only in the extent of activation in these regions. Neither group replicates the substantial right hemisphere activation found in the picture naming task. Cargnelutti et al.’s (2019) AoA analyses present a similar picture: the clusters found for early bilinguals are subsumed within the larger scope of activation generated in the late bilinguals. For the L1 network, neither AoA group overlaps with the picture naming task. In the L2 network, both AoA groups replicates clusters in the left *inferior frontal gyrus* (early BA 47, late BA 9) and *medial frontal gyrus* (BA 6), with an additional cluster in the *insula* replicated by late bilinguals. Some of the findings appear to switch from the L1 to L2 network between analysis, for example, posterior activation in the right hemisphere is greater for the L2 of late bilinguals and the L1 in the picture naming task.

Therefore, in considering what effect participant characteristics may have on the analysis, it appears that during picture naming tasks, <u>low proficiency bilinguals</u> engage the left posterior temporal lobe in their **L1 network**. In particular, early bilinguals rely more upon posterior regions, and late bilinguals recruit greater neural resources across these regions. The **L2 network** of low proficiency bilinguals involves bilateral frontal and subcortical activation, largely in the left *cingulate* and *paracingulate gyrus*. <u>High proficiency bilinguals</u> also engage a slightly different set of left posterior regions in the **L1 network**, and left frontal regions in the **L2 network** for both early and late bilinguals. However, late bilinguals produce a broader network with bilateral subcortical activation, particularly in the right *caudate* and bilateral *insulae*. Early bilinguals generate a core region within this broader L2 network. The widespread right hemisphere activation observed in the task analysis for the **L1 network** may be unique to picture naming more generally, as this finding did not replicate in earlier analyses. The finding could also be an artifact of the data set; however, the picture naming task is one of the more highly powered analyses for both networks, and the task is simple enough not to allow for a huge amount of variation. All but two foci for this analysis were taken from experiments that utilized line drawings as stimuli.

### 4.3 Passive Listening

The previous meta-analyses that differentiate linguistic competencies also lack a category that represents passive listening tasks. Indefrey’s (2006) sentence-level listening and reading category included six passive listening tasks out of fourteen experiments, although only two of these experiments contributed to the findings. Nonetheless, two of the three clusters obtained in the L2 network of the earlier study replicate in the task analysis in the *middle frontal gyrus* (BA 9) and the *inferior frontal gyrus* (BA 44). Additional activation in the bilateral *superior temporal gyrus* (left BA 41, right BA 22) and left *caudate body* were not reflected in Indefrey’s (2006) more expansive category. In Sulpizio et al.’s (2020) meta-analysis, passive listening tasks were divided among the linguistic categories, with just one passive listening task included in each contrast. An exception can be found in the phonology domain for the L1 network, in which three of the five tasks involved passive listening. In the L2 network, only one passive listening task was present. Therefore, only the results of the L1 network will be compared to our passive listening task. The earlier meta-analysis found one cluster in the left *inferior frontal gyrus*, which did not replicate in the passive listening task. In the right hemisphere, activation in the *middle temporal gyrus* (BA 21/22) was partially replicated in the right *sub-gyral temporal lobe* (BA 21) in the task analysis. The fact that the earlier study supports this finding in the L1 network despite the limited data set available for the passive listening task may justify the validity of our assumption that there should be high confidence in our data selection process, such that even smaller data sets are reliable despite their limited size.

Overall, there is little prior data to support a comparison between previous meta-analyses and the passive listening analysis, particularly for the L2 network. However, the results that are available generally confirm our findings. We propose that the bilateral temporal activation found in the task analysis may be specific to passive listening. The L1 network described in the task analysis is generally supported by the most relevant previous meta-analysis. The fact that our data set is more specific to the task of interest than the phonology data set would suggest that activation in frontal regions found on the phonology domain is unlikely to be a consistent feature of the L1 network for passive listening. The passive listening task engaged roughly half of the regions identified as the language control network by Luk et al. (2012), but only in the L2 network and primarily in the left hemisphere. These regions included the left *middle frontal gyrus* (BA 9), *inferior frontal gyrus* (BA 44), *caudate*, and right *superior temporal gyrus* (BA 6). The *precentral gyrus* (BA 6) and *midline pre-supplementary motor area* (BA 6) are reported in Luk et al. (2012), whereas passive listening engages this area in the *superior fontal gyrus* (BA 6).

When considering proficiency level, the studies that contributed foci to the task analysis predominately involved moderate to high proficiency bilinguals. A single study with low proficiency bilinguals did contribute over half of the foci used in the L2 analysis, but generated few of the resulting clusters. Nonetheless, Sebastian et al.’s (2011) low proficiency analysis more closely approximates the results of the L1 network in the task analysis, replicating the bilateral activation in the *middle temporal lobe* (BA 21), whereas no common activation was found for the high proficiency group. In the L2 network, the passive listening task again shows no overlap with the high proficiency analysis, although one cluster was found by Sebastian et al. (2011) in a more anterior region of the right temporal lobe. Despite its broader scope of activation, the low proficiency analysis replicates just one cluster found in the task analysis in the *middle frontal gyrus* (BA 9), with an adjacent cluster in the *inferior frontal gyrus* (BA 44). Cargnelutti et al.’s (2019) analysis of highly proficient bilinguals show no overlapping activation in the L1 network for the late bilinguals, and a shared cluster in the left *middle temporal gyrus* (BA 21) for early bilinguals. In the L2 network, a commonly activated region in the classical language network, the left *inferior frontal gyrus* (BA 9/44) is replicated across studies, for both the late and early bilinguals.

When considering AoA, all but one of the studies that contributed foci to the task analysis involved participants with moderate AoA. Liu & Cao’s (2016) AoA analyses poorly represent the passive listening results, suggesting that AoA is not particularly meaningful for distinguishing L1 and L2 networks. In the L1 network, early and late bilinguals generate one cluster in the left *superior temporal gyrus* (early BA 22, late BA 22/38) that is adjacent to activation in the task analysis. In the L2 network, early bilinguals generate clusters adjacent to the left *middle* and *inferior frontal gyrus* (BA 9) and *superior frontal gyrus* (BA 6). Late bilinguals replicate clusters found in the task analysis in the left *middle frontal gyrus* (BA 9), *superior frontal gyrus* (BA 6), and an adjacent cluster in the *inferior frontal gyrus* (BA 44). Cargnelutti et al.’s (2019) high proficiency analysis also poorly represents the passive listening task. In the L1 network, early bilinguals replicate a cluster in the left *middle temporal gyrus* (BA 21). In the L2 network, both groups replicate a cluster in the left *inferior frontal gyrus* (early BA 44, late BA 9/44). Late bilinguals reproduce additional clusters from the task analysis in the left *superior temporal gyrus* (BA 41) and caudate. None of the posterior activation found in the AoA analysis is observed in the task analysis.

Therefore, in considering what effect participant characteristics may have on the task analysis, it appears that during passive listening tasks, <u>low proficiency bilinguals</u> engage bilateral temporal regions associated with sound processing in the **L1 network**. This is true for early and late bilinguals, although late bilinguals may recruit more neural resources in the left hemisphere. The **L2 network** of low proficiency bilinguals engages frontal regions of the left hemisphere. <u>High proficiency bilinguals</u> activate the left *inferior frontal gyrus* for both early and late bilinguals, whereas only late bilinguals engage left temporal regions. Previous meta-analysis do not account for one finding from the passive listening analysis: substantial right hemisphere activation of the *superior temporal gyrus*. It may be that this finding is specific to the task analysis and therefore was not identified in early studies. It is also important to note that stimuli from the listening tasks utilized in the L1 analysis were primarily on the discourse level, and on the sentence level for the L2 analysis. It may be that more complex listening stimuli elicit greater engagement of temporal regions, but this will require additional study given the limited L1 data set.

### 4.4 Syntactic Judgment

Syntactic judgment is not included among the linguistic competencies described by Indefrey (2006), although one experiment from the syntactic judgment task is found in the sentence-level listening and reading data set. This study contributed to activation in the L2 network in the left *middle frontal gyrus* (BA 9) and *inferior frontal gyrus* (BA 44, BA 47). The task analysis replicated clusters in the *middle frontal gyrus* (BA 9) and *inferior frontal gyrus* (BA 44). However, the task analysis generated a substantially wider response in left motor and posterior regions, as well as in bilateral subcortical areas. In Sulpizio et al.’s (2020) grammar domain, roughly half of the experiments in the data set contain syntactic judgment tasks. In the L1 network, the earlier meta-analysis finds numerous clusters in traditional language areas. However, the task analysis replicates just one cluster in the bilateral *insulae* (BA 13). Two additional clusters in the right hemisphere are located adjacent to activation from the earlier study, in the *medial frontal gyrus* (BA 6) and *fusiform gyrus* (BA 20). In the L2 network, the task analysis again generates substantially more activation. Clusters in the left *inferior frontal gyrus* (BA 44) and *putamen* replicated, and a cluster adjacent to the earlier analysis is found in the left *precuneus* (BA 19). Discrepancies between the two studies concern the scope of activation for each language, with the grammar analysis generating a larger L1 network and the task analysis a larger L2 network. The smaller network in each study that engaged the *cerebellum*.

Overall, there is a lack of consistency between the current analysis and previous ones, particularly regarding engagement of core language areas. The previous meta-analyses included few true syntactic judgment tasks, and therefore the syntactic judgment task is unlikely to be defined by strong engagement of traditional language areas, particularly the left *inferior frontal gyrus*. The L1 network shows some consistency between analyses, but the task analysis generates greater right hemisphere involvement, with right frontal and temporal activation, as well as bilateral subcortical regions. The cerebellum is activated in this network, which may reflect procedural memory attributed to automatic recall. These findings are consistent with observations made about L1 processing more generally. In the more extensive L2 network, left frontal activation was found, with additional clusters in posterior regions and the *putamen*. This frontal and subcortical activation may be bilateral. However, much of the frontal activation is outside of language areas, localized in superior regions or in motor areas. The language control network was engage in the left hemisphere of the L2 network in the *middle frontal gyrus* (BA 9), *inferior frontal gyrus* (BA 44), and *caudate*, as well as in the regions adjacent to those identified in Luk et al. (2012): the left *fusiform gyrus* (BA 37) and *precentral gyrus* (BA 6).

When considering proficiency level, the syntactic judgment data set for the L1 network was composed entirely of studies with highly proficient bilinguals, as were the majority of experiments for the L2 network. Sebastian et al.’s (2011) high proficiency group generates activation in left posterior regions in the L1 network, whereas the results of the task analysis are almost entirely subcortical or right-lateralized, leading to no shared clusters in either hemisphere. In the L2 network, an adjacent cluster is found in the task analysis in the left *middle frontal gyrus* (BA 9). Low proficiency bilinguals generated no shared activation in the L1 network. In the L2 network, the task analysis replicated a cluster in left *middle frontal gyrus* (BA 9) and produced an adjacent cluster in the left *inferior frontal gyrus* (BA 44) and *precuneus* (BA 19). Each study generates substantial subcortical activation, but in different regions. Cargnelutti et al.’s (2019) high proficiency analysis also generated no shared activation for early or late bilinguals in the L1 network. In the L2 network, the left *inferior frontal gyrus* (BA 9/44) replicated for both groups, and late bilinguals produced activation that was adjacent to the task analysis in the left *inferior parietal lobule* (BA 40) and bilateral *caudate*.

When considering AoA, the syntactic judgment data set for the L1 network largely represents bilinguals with moderate AoA, and the L2 network contains a balance of late and moderate bilinguals. In each network, just one study includes early bilinguals. In Liu & Cao (2016), early and late bilinguals show substantial overlap with one another in both networks. In the L1 network, early and late bilinguals produced clusters that are adjacent to the task analysis in the left *postcentral gyrus* (BA 40). In the L2 network, early bilinguals replicated activation in the task analysis in the left *inferior frontal gyrus* (BA 9) and produced an adjacent cluster in the left *middle frontal gyrus* (BA 9). Late bilinguals reversed this order, replicating a cluster in the left *middle frontal gyrus* (BA 9) and generating an adjacent cluster in the left *inferior frontal gyrus* (BA 44) and *precentral gyrus* (BA 6). Cargnelutti et al. (2019)’s AoA analysis for the L1 network of late bilinguals replicated a cluster in the left *insula* (BA 13) and found an adjacent cluster in the right *medial frontal gyrus* (BA 6). No common clusters were found for early bilinguals. In the L2 network, the early and late bilinguals replicated the left *inferior frontal gyrus* (BA 9/44). Early bilinguals replicated the left *precentral gyrus* (BA 6) and late bilinguals replicated the left *inferior parietal lobule* (BA 40). More extensive frontal activation and a lack of temporal activation were key differences between the earlier findings and the results of the syntactic judgment task. Additionally, involvement of the bilateral *insulae* was found for late bilinguals in the L2 network in the earlier study, but in the L1 network for syntactic judgment.

Therefore, in considering what effect participant characteristics may have on the task analysis, it appears that during syntactic judgment tasks, <u>low proficiency bilinguals</u> show minimal to no activation of the **L1 network**. Potentially, late bilinguals may engage frontal and subcortical regions, and early bilinguals may involve the left *insula* and right motor areas. In the **L2 network**, early and late bilinguals activated frontal regions, but late bilinguals did so more substantially and engaged posterior regions to a lesser degree. <u>High proficiency bilinguals</u> may engage left posterior regions in the **L1 network**, with some evidence of motor and subcortical involvement in late bilinguals. In the **L2 network**, left frontal regions were engaged for all bilinguals, whereas inclusion of posterior and subcortical activation was evident in late bilinguals. The specific location and extent of subcortical activation appears specific to the syntactic judgment task, as does the activation of BA 9 for the L2 across groups.

It is important to note that all the results for the L1 network came from one study, Saur et al. (2009), indicating the limited scope of this data set, and therefore the degree of confidence that should be attributed to those results. Saur et al. (2009) was the only study in the task to present stimuli aurally. In the L2 network, experiments conducted by Sakai et al. (2009) had the greatest weight, yet contributed just one focus. All other studies in the data set were relatively balanced in their weight, suggesting a more consistent outcome for the L2 analysis.

### 4.5 Semantic Judgment

The semantic judgment task corresponds to the semantic decision category described by Indefrey (2006). This category yielded one cluster in the L1 network in the left *anterior middle temporal gyrus* (BA 21). The semantic judgment task generated a cluster in the same region in the right hemisphere and an adjacent cluster in the left *superior temporal gyrus* (BA 39). The L2 network of the semantic judgment task replicates a cluster in the left *middle frontal gyrus* (BA 9), with adjacent cluster in not the anterior but *posterior cingulate* (BA 29). Another prominent cluster is found in an anatomical region common to both analyses (BA 39). In Sulpizio et al. (2020), only a small number of studies in the lexico-semantic domain consist of semantic judgment tasks: three of eighteen in the L1 data set and two of twenty-two studies in the L2 data set. No activation replicates in the L1 network, and only the left *prefrontal gyrus* (BA 6) and right *insula* (BA 13) replicate in the L2 network. The frontal activation found in both networks of the lexico-semantic analysis is notably absent in the task analysis.

Overall, the results observed for the semantic decision task bear a striking similarity to the results of the syntactic decision task. One study (three experiments) is shared between their data sets; however, this study was not among the most heavily weighted in either task analysis and contributed no foci to their L1 networks. Thus, the two decision tasks independently produced similar results across different sets of experiments. Shared aspects of task activation include motor areas such as the bilateral *precentral* or *medial frontal gyrus* (BA 6) for both networks and engagement of the cerebellum instead of traditional language areas, particularly the left *inferior frontal gyrus*. Differences can be found in the location of posterior activation, which may be specific to each linguistic competency instead of the decision component of the task. Posterior clusters included the *middle temporal gyrus* (BA 39, semantic processing) for the semantic task versus the *fusiform gyrus* in the syntactic task (BA 37, morphological processing). Both tasks engaged left subcortical regions: the *posterior cingulate* for semantic tasks and the *putamen* for syntactic tasks. In the L1 network, the tasks also generated similar right hemisphere activation, especially in the *precentral gyrus* (BA 6). Differences in this network that may be task-specific are engagement of the *superior temporal gyrus* (BA 39) for semantic judgment, and the *postcentral gyrus* and *cerebellum* for syntactic judgment.

A number of regions in both networks belong to the language control network. In the L1 network, this is the left *precentral gyrus* (BA 6). We find bilateral activation in the *temporal lobe* (left BA 39, right BA 21), although the precise location reverses the areas identified by Luk et al. (2012) between hemispheres. A cluster in the *precentral gyrus* (BA 6) of the L2 network in the opposite (right) hemisphere may also be relevant. The L2 network activates the left *middle frontal gyrus* (BA 9) and the *middle temporal gyrus* in an anatomically adjacent area (BA 39), as well as the right *caudate*. These regions are similar to those engaged in the L2 network of the syntactic decision task, although the syntactic task arguably involves the full range of language control regions, with the additional activation of the *inferior frontal gyrus* (BA 44) and bilateral *caudate*.

When considering proficiency level, the semantic judgment data set for both networks was evenly balanced between high and low proficiency levels. Sebastian et al.’s (2011) high proficiency analysis finds no common activation in the L1 network. In the L2 network, one adjacent cluster in the *middle frontal gyrus* (BA 9) was identified. Low proficiency bilinguals replicated a cluster in the L1 network of the task analysis in the right *middle temporal gyrus* (BA 21) and produced an adjacent cluster in the left *superior temporal gyrus* (BA 39). In the L2 network, the earlier analysis reproduces a clusters from the task analysis in the left *middle frontal gyrus* (BA 9) and *precuneus* (BA 7). The analysis generates adjacent clusters in the left *cingulate gyrus* (BA 23) and *precentral gyrus* (BA 6). In the right hemisphere, the claustrum replicates. Cargnelutti et al.’s (2019) high proficiency analysis also generated no shared activation with the L1 network for either early or late bilinguals. In the L2 network, this was also true of early bilinguals. Late bilinguals replicated a cluster from the task analysis in the left *middle temporal gyrus* (BA 39) and produced an adjacent cluster in the left *angular gyrus* (BA 39).

When considering AoA, the semantic judgment data set largely represents bilinguals of a moderate or late AoA for both networks. Liu & Cao’s (2016) AoA analyses poorly represent the L1 network observed in this task, identifying just one cluster for early and late bilinguals that was adjacent to the task analysis in the left *superior temporal gyrus* (BA 39). In the L2 network, early bilinguals replicated a cluster found in the task analysis in the left *precentral gyrus* (BA 6) and produced an adjacent cluster in the left *middle frontal gyrus* (BA 9). These findings were reversed in late bilinguals, who replicated the second cluster (BA 9) and found a cluster adjacent to the first (BA 6). Cargnelutti et al.’s (2019) AoA analysis finds no common activation in the L1 network for either early or late bilinguals, with the exception of an adjacent cluster found in both groups in the left *superior temporal gyrus* (BA 39). In the L2 network, early bilinguals replicated a cluster from the task analysis in the left *precentral gyrus* (BA 6). Early and late bilinguals both generated a cluster adjacent to the left *middle temporal gyrus* (BA 39) and *angular gyrus* (BA 39). Late bilinguals replicated a cluster in the right *insula* (BA 13) and produced a second adjacent cluster in the *precuneus* (BA 7).

Therefore, in considering what effect participant characteristics may have on the task analysis, it appears that during syntactic judgment tasks, <u>low proficiency bilinguals</u> produce bilateral temporal activation in the **L1 network** for all bilinguals. This activation may be more extensive and located in semantic processing areas of the left hemisphere and in sound processing areas of the right hemisphere. In the **L2 network**, a broad network is engaged in frontal and posterior regions in the left hemisphere by late bilinguals. This involves left frontal, motor, and posterior regions, as well as bilateral activation of subcortical regions. The frontal and motor regions are jointly activated for early and late bilinguals. Early bilinguals may engage BA 6 more strongly, whereas late bilinguals may tend to engage posterior areas. <u>High proficiency bilinguals</u> show no common activation in the **L1 network**. In the **L2 network**, all bilinguals involve frontal regions outside of the classical language network in the left hemisphere. Early bilinguals may involve left motor regions, and late bilinguals may engage right subcortical areas and greater left posterior regions. This decision task also engaged BA 9 across all L2 groups.

## 5. Conclusion

Neuroimaging studies that offer clear and unambiguous answers to a falsifiable hypothesis are often considered best practice and the most convincing or useful contributions to the field. However, it can be seen from a review of previous meta-analyses that attempting to create simplicity by neglecting some inherent complexity in the problem can lead to results that raise additional questions or do not converge with other research. When faced with this dilemma in the bilingual neuroimaging literature, this paper proposes adopting the approach of triangulating data, to see what replicates across analyses, each of which individually may have less power or be partially representative of the underlying population. Instead of creating the largest meta-analysis by consolidating across numerous factors, we have conducted five smaller analyses that can be compared to one another and to earlier meta-analyses that have been designed to address their own research questions.

Some general assumptions of how common factors affect L1 and L2 processing appear to hold true across tasks. This includes the effect of AoA and proficiency level, such as the tendency for high proficiency bilinguals to engage more frontal regions and for low proficiency bilinguals to involve more temporal and posterior regions, which has sometimes been interpreted as a shallower degree of language processing. Across most tasks, the L2 network recruited a wider and more bilateral network, and right hemisphere activation appeared to be influenced more strongly by proficiency level than AoA. However, several unanticipated findings were revealed. In particular, although previous literature has raised the question of greater subcortical activation arising in one language network (cf. Hernandez and Li, 2007), the degree to which subcortical activation varied by task, in terms of the extent, location, and language network that engaged these subcortical regions was surprising and requires additional investigation. The greatest degree of subcortical activation was elicited by tasks that required an active categorical response, such as a button-press, or the attribution of the correct name to a picture.

When we look at task-specific variation, considerable similarity was found between the two decision tasks, both of which appeared to rely not upon frontal regions for task-specific processing, but instead strongly engaged motor regions and the language control network. Their L1 network was minimally engaged, particularly for low proficiency bilinguals, yet it was the L1 networks, rather than more extensive L2 networks that diverged most substantially, supporting the idea that there may be a fundamental difference in the way bilinguals process their first and second language. In fact, in regards to the subcortical activation, it was the picture naming task that more closely resembled the results of the semantic decision task. Another difference between the results of the decision tasks may be motivated either by the nature of the task, or by the experimental design. The syntactic stimuli were largely sentence-level items, whereas the majority of semantic tasks, including the most heavily weighted studies, utilized word-level stimuli. The syntactic task engaged morphological processing areas, and the semantic task involved bilateral temporal regions involved in processing word meaning.

Other partial similarities were found across tasks, in either their L1 or L2 network, but rarely in both, again suggesting that the two networks may be independent. Bilateral temporal activation was exhibited by the semantic judgment task and the passive listening task, but the first engaged word meaning areas in the left hemisphere and sound processing areas in the right hemisphere, whereas passive listening involved only sound processing areas in either hemisphere. In the L2 network, bilateral temporal regions were also engaged, but higher up in the *superior temporal gyrus*. The L1 network of the reading and picture naming tasks both incorporated widespread right hemisphere activation; however, the reading task was more balanced bilaterally, with clusters in the L1 and L2 network appearing in close proximity, whereas the L1 network of the picture naming task was clearly right-lateralized. Additionally, the reading task activated all of the core language processing areas in both the L1 and L2 network. In the picture naming task, the L1 and L2 networks diverged dramatically, with the L1 network clearly dominated by posterior regions and the L2 network by frontal regions. These two tasks were also the only two to show substantial frontal activation in their L2 network, perhaps in conjunction with the complex nature of their task paradigms.

It is these intersecting similarities and differences that make a case for the unique composition of task-based and language-based networks, just as the confirmation of widely held assumptions about the effect of secondary factors on linguistic processing provides some legitimacy for the analysis, despite the smaller size of the corpus. However, even a careful inter pretation of the data is still limited by the shortcomings of the data set, which at this point cannot fully represent the full complexity of bilingual language processing and does not meet the highest level of standards set by the originators of the ALE method to ensure well-powered studies. The aim of this paper is to foreground the inconsistencies that currently exist among meta-analyses of the bilingual neuroimaging literature, to provide some compelling hypotheses for further research, and to clarify best practices and what steps are needed to ensure that the data necessary for this type of research will become available to the wider research community in the future.

